# An ecological perspective on microbial genes of unknown function in soil

**DOI:** 10.1101/2021.12.02.470747

**Authors:** Hannah Holland-Moritz, Chiara Vanni, Antonio Fernandez-Guerra, Andrew Bissett, Noah Fierer

**Affiliations:** Department of Natural Resources and the Environment, University of New Hampshire, Durham, NH, USA; Microbial Genomics and Bioinformatics Research Group, Max Planck Institute for Marine Microbiology, Celsiusstraße 1, 28359, Bremen, Germany, Jacobs University Bremen, Campus Ring 1, 28759 Bremen, Germany; Lundbeck GeoGenetics Centre, The Globe Institute, University of Copenhagen, Copenhagen, Denmark; CSIRO, Oceans and Atmosphere, Hobart, Tasmania, Australia; Department of Ecology and Evolutionary Biology, Cooperative Inst. for Research in Environmental Sciences, University of Colorado, Boulder, USA

## Abstract

Genes that remain hypothetical, uncharacterized, and unannotated comprise a substantial portion of metagenomic datasets and are likely to be particularly prevalent in soils where poorly characterized taxa predominate. Documenting the prevalence, distribution, and potential roles of these genes of unknown function is an important first step to understanding their functional contributions in soil communities. We identified genes of unknown function from 50 soil metagenomes and analyzed their environmental distributions and ecological associations. We found that genes of unknown function are prevalent in soils, particularly fine-textured, higher pH soils that harbor greater abundances of *Crenarchaeota, Gemmatimonadota, Nitrospirota*, and *Methylomirabilota*. We identified 43 dominant (abundant and ubiquitous) gene clusters of unknown function and determined their associations with soil microbial phyla and other “known” genes. We found that these dominant unknown genes were commonly associated with microbial phyla that are relatively uncharacterized, with the majority of these dominant unknown genes associated with mobile genetic elements. This work demonstrates a strategy for investigating genes of unknown function in soils, emphasizes the biological insights that can be learned by adopting this strategy, and highlights specific hypotheses that warrant further investigation regarding the functional roles of abundant and ubiquitous genes of unknown function in soil metagenomes.

## Introduction

Soils are one of the largest reservoirs of genetic diversity on Earth (1,2). One gram of soil can harbor hundreds to thousands of distinct microbial taxa (3,4) and an estimated 1 000 Gbp of microbial genome sequences (5). Soil microbial communities are essential contributors to terrestrial nutrient cycling and carbon dynamics, with wide-ranging effects on plant and animal health (6). Unfortunately, soil microbial taxa are underrepresented in culture and genomic sequence databases (6,7). The adoption of high-throughput sequencing technologies has yielded new insights into the full-extent of soil microbial diversity and provide glimpses into the genomes, transcriptomes, and functional attributes of organisms that are as-yet uncharacterized. While these molecular approaches bring clear benefits to understanding soil microbial communities, they also pose a fundamental challenge as a large fraction of genomic data is composed of genes of unknown function.

Genes of unknown function are abundant in many habitats (40-60% of genes in a given environment (8–11)). While this issue is not unique to soils, several aspects of soil microbiology indicate that soil microbial communities harbor large proportions of genes of unknown function. First, soils are one of the most phylogenetically diverse microbial habitats (12,13) and the genetic diversity found within soils is equally impressive (2,14,15). Second, the majority of soil microbes are difficult to isolate and study in the laboratory (16). There are many reasons for this, including the observation that many soil microbes are likely slow-growing oligotrophs that can be difficult to culture using standard approaches (17). Thus, soil environments often harbor a higher proportion of uncultivated taxa (18,19) as compared to environments (like human skin and gut) where a larger fraction of the microbial communities are represented in culture-based collections (16). Third, many physical and biological aspects of soil, such as micro-site heterogeneity and high strain-level variation, make it a particularly challenging environment to apply assembly-based metagenomic techniques (6,20). Despite these limitations, the genes of unknown function found in soil may hold important ecological information (9,21). From work with marine and human microbiomes (11), we know that these genes are enriched in phyla with more genomes represented only in metagenome-assembled genomes (MAGs), have more restricted distributions than known genes (i.e., less likely to be shared across distinct samples), and are more common in marine environments than in host-associated environments. Metagenomic studies of soil microbial communities typically focus only on “known” genes, i.e. genes that can be functionally annotated with a reasonably high degree of confidence (with some notable exceptions, e.g. (19)), effectively discarding the unannotated genes from downstream analyses and ignoring the insight that these genes may provide about microbial life in soil. While these unannotated genes are often acknowledged (2,11,12), they are typically not the explicit focus of study.

We set out to understand the genes of unknown function in 50 soil samples from a publicly available dataset compiled by the Australian Microbiome Initiative (22). Specifically, we used a new tool, AGNOSTOS (11), to identify gene clusters and group them into four categories: “Known with Pfam Annotation”, “Known without Pfam Annotation”, “Genomic Unknowns” (found in cultured and sequenced isolates and in MAGs of sufficient quality to be included in genome databases), and “Environmental Unknowns” (found in metagenomes) – see Table 1. Using these data, we asked the following questions: 1) How abundant are genes of unknown function? 2) Are there particular soil conditions, or soils containing particular microbial taxa, that are more likely to harbor higher abundances of these genes? 3) Which genes of unknown function are particularly abundant and ubiquitous across soils? Broadly, we hypothesized that genes of unknown function would be widespread in soils and the proportion of these genes found in metagenomes would vary along ecological gradients, with higher proportions in soils where environmental conditions limit the abundances of more copiotrophic taxa (19). We further hypothesized that the genes of unknown function would be enriched in soil that contained higher abundances of under-studied microbial lineages (i.e. lineages with few cultivated representatives). Finally, we compiled a “hit list” of abundant and ubiquitous (“dominant”) genes of unknown function in soils and used their associations with microbial taxa and genes of known function to infer their potential ecological characteristics and direct future research efforts to further explore the novel genetic diversity found in soil microbial communities.

**Table 1:** Description of each gene cluster category.

| Category | Description |
| --- | --- |
| **Knowns** |  |
| Known with Pfam annotation | Gene cluster contains at least one Pfam domain of known function |
| Known without Pfam annotation | Gene cluster contains genes that are without Pfam domain of known function, but are annotated with a function in the NCBI *nr* or UniRef90 databases |
| **Unknowns** |  |
| Genomic unknown | Gene cluster contains at least one gene that is found in sequenced and draft genomes but are hypothetical, uncharacterized, or otherwise unannotated |
| Environmental unknown | Gene cluster contains genes that have no known sequence homology in sequenced or draft genomes and only found in metagenomes and MAGs |

## Methods

### Sample selection

We used publicly available soil metagenomic data from the Australian Microbiome Initiative Biomes of Australian Soil Environments (BASE) project (22). The project includes 449 soil samples for which shotgun metagenomic data are available. From this subset of samples that had metagenomic data, we selected a random sub-sample of 48 surface soil samples and viewed them on a map to make sure that they represented a good spatial sampling of regions across Australia (Figure S1A). The goal of this random sub-sampling was to reduce the number of samples to a computationally manageable size and to select a set of samples that represented a wide range of environmental gradients across Australia. The sample identifiers are listed in Table S1 and raw sequencing data are available at the Australian Microbiome Initiative Data Portal (https://data.bioplatforms.com/organization/about/australian-microbiome) as well as on the NCBI Sequence Read Archive (BioProject accession number PRJNA317932).

### Sampling protocol, DNA extraction, and sequencing

The Australian Microbiome Initiative uses a standard set of protocols for all samples. Full details are available at https://www.australianmicrobiome.com/protocols/. Specific protocols used for the samples and data included in this study are found under protocol headings labeled (“as per BASE”). To summarize, between 9-30 soil samples were taken from the top 10 cm of a 25×25 m plot with the plot location selected to represent a visually homogeneous soil environment. Soil samples from the plot were then homogenized to generate one representative surface sample per plot with the composited sample frozen immediately after collection.

DNA was extracted from triplicate 250 mg sub-samples of each soil using the DNeasy PowerLyzer PowerSoil Kit (Qiagen), following the Earth Microbiome protocol (23), and pooled prior to downstream processing. Metagenomic sequencing libraries were prepared following the Illumina TruSeqNano DNA Sample Preparation Guide. Briefly, 200 ng of DNA from each sample was sheared to 550bp using Covaris sonic shearing. Clean-up of the sheared DNA, ligation of the adapters, and PCR amplification of the DNA followed the TruSeq Nano DNA Sample Preparation Guide. The resulting libraries were sequenced on an Illumina HiSeq with 2×150bp sequencing chemistry. The median number of reads per sample was 30.2 million reads, with a range from 11.8 million reads to 154.9 million reads (Table S1).

### Environmental data

To better understand the relationships between genes of unknown function and environmental gradients, we selected ten soil and site factors that are often important in structuring soil microbial communities (6). The factors included four site-specific variables: mean annual temperature, mean annual precipitation, aridity (as inferred from calculation of the aridity index, (24)), and net aboveground primary productivity (NPP). We also included six soil-specific variables: soil pH, percent organic carbon, extractable inorganic nitrogen, phosphorus, conductivity, and texture (% silt + % clay) (Figure S1B). We calculated Pearson correlations between each of these environmental variables. Most were weakly to moderately correlated in our samples, except site aridity index and mean annual precipitation, which were understandably well-correlated as precipitation is used to calculate the aridity index (Figure S1B). As is evident from Figure S1C, the 48 samples span broad gradients in soil and site characteristics (Figure S1C).

Analyses of soil chemical and physical properties were conducted by a commercial laboratory (CSBP Laboratories, Bibra Lake, Western Australia) using standard methods. Briefly, percent organic carbon was measured according to the Walkley-Black method using concentrated sulfuric acid and dichromate in a colorimetric test. For extractable inorganic nitrogen, we summed extractable nitrate and ammonium together. Both nitrate and ammonium were extracted with 2M KCl and measured colorimetrically. Available phosphorous was calculated using the Colwell method (25). We also summarized the texture of each soil into a single metric by summing the silt and clay percentages.

For each soil, we also compiled data on climate and plant productivity (i.e., mean annual temperature, mean annual precipitation, aridity index, and NPP) at the collection site from the Atlas of Living Australia spatial portal (26). From this data portal, we selected the mean annual precipitation, aridity index, and NPP standard environmental data products produced by the ecosystem sciences division of Australia’s Commonwealth Scientific and Industrial Research Organisation (CSIRO), and the mean annual temperature product from the WorldClim database (27).

### Generation of metagenomic assemblies and taxonomic composition table

We downloaded the raw sequence data from the Bioplatforms Australia Data Portal (https://data.bioplatforms.com/). After downloading, we removed adapters and other library-preparation contaminant sequences from the FASTQ files using Cutadapt (28) and filtered out sequences with low average quality scores and short sequence length with Sickle (29) (-q 20 -l 50). We verified the read quality of our filtered data with FastQC (http://www.bioinformatics.babraham.ac.uk/projects/fastqc/) and MultiQC (30) before proceeding. To assemble the reads, we tried three different assemblers (IDBA – settings: default (31), metaSPAdes – settings: default (32), and MEGAHIT – settings: “meta-large” preset (33)) and ultimately used MEGAHIT as it represented a good balance between assembly quality and computational burden. To minimize downstream computational costs, we filtered out small contigs <1000 bp, since these were unlikely to contain whole genes. We assessed assembly quality with QUAST (default settings) (34), and used Bowtie2 (default settings) (35) to map the filtered reads back to the filtered assemblies.

As is common for soils, assembly rates were fairly low with an average assembly length of 123 Mbp (ranging from 7 – 954 Mbp), and a mean N50 of 0.79 Kbp (ranging from 0.6 – 1.3 Kbp) before filtering. We note that, although the assemblies are fragmented, our analyses focused on dominant and abundant gene clusters (see below), which are less likely to be affected by the low assembly rates. Assembly, rather than read-based analysis, is required for downstream analysis with AGNOSTOS (see below). Full assembly statistics and mapping rates can be found in Table S1.

We also determined the taxonomic composition of the microbial communities in each sample. We used phyloFlash (36) in “almost everything” mode to identify and assign taxonomy to fragments of the small-subunit (16S/18S) rRNA gene in each metagenome. Using the number of hits to fragments identified in each sample, we created a taxon-by-sample table for downstream analyses. After creating the table, we filtered the table to exclude taxa identified as Chloroplast, Mitochondria, or Eukaryotes, with these categories typically representing <7.2% of recovered small-subunit rRNA gene reads. We also removed any taxa that were unassigned at the phylum level (<3.5% of small-subunit rRNA gene reads). After filtering the table, we controlled for differences in sequencing depth by rarefying the table. We picked 1720 rRNA gene hits at random from each sample, which was the number of hits in the sample with the lowest number of hits. Finally, we converted our taxon-by-sample table into proportional abundances for downstream analyses.

### Identifying genes of unknown function

To identify genes of unknown function in our samples we used AGNOSTOS (11). AGNOSTOS is a tool for partitioning coding sequence space into known and unknown clusters of genes. It clusters genes based on sequence homology and classifies each gene cluster into one of four categories, *Known with Pfam Annotation, Known without Pfam Annotation, Genomic Unknown*, and *Environmental Unknown* (Table 1). To integrate soil metagenomes with the original AGNOSTOS database (seedDB+NCLDV), we used the “DB-update” module of AGNOSTOS described in (37) which integrates new sequences into the original gene cluster database. We note that the AGNOSTOS database is dominated by prokaryotic (bacterial and archaeal) genes, as genes from eukaryotes (including fungi) and viruses represent only a small fraction of the total database (37). Briefly, AGNOSTOS uses Prodigal (38) to identify open reading frames in assembled contigs and filters out spurious open reading frames that overlap with true protein-coding open reading frames (commonly called “shadow genes” (38) and those that are found in the Antifam database of spurious open reading frames (40). Importantly for metagenomic analysis, partial and incomplete genes are retained. AGNOSTOS then annotates these genes against Pfam (41) using hmmsearch from HMMER (42). After the Pfam annotation, genes are clustered with MMseqs2 (43) to create a homology-based database that is independent of a gene cluster’s known or unknown status. AGNOSTOS then screens the gene clusters for quality, ensuring that each cluster is made up of sufficiently homologous genes, and discarding those clusters composed of insufficiently similar genes and those composed of a high number of spurious genes. At this stage, clusters containing singletons are identified, but were not further categorized as there is insufficient information in a singleton cluster for reliable categorization into one of the four categories (Table 1). Although singleton clusters lack sufficient information to rigorously categorize, it is likely that the majority of these clusters represent genes of unknown function that are rarer in databases and in our samples. We note that Vanni and others (37) found that, as new data is added to the AGNOSTOS database, many of the singleton gene clusters acquire more representation and can be identified and characterized, suggesting that the majority of these sequences are not sequencing artifacts or assembly errors, but simply rarer genes. After clusters are cleaned and filtered (including removing singletons), a consensus sequence for each cluster is generated and clusters are each assigned to one of the four categories (Table 1). Gene clusters with at least one member assigned to a Pfam domain of known function are classified as “known with Pfam”. Gene clusters with a consensus sequence that matched sequences in the NCBI *nr* database (44), UniRef90 database (45) or the Uniclust database (46), are categorized as either “known without Pfam annotation” or “genomic unknowns” depending on their similarity to characterized proteins or to proteins annotated with terms typical of genes of unknown function (e.g. “hypothetical”, “uncharacterized”, etc.) or another name used to classify unknown proteins in the databases’ controlled vocabulary. Gene clusters with Pfam annotations to domains of unknown function (“DUF”) are then also added in the “genomic unknown” category. Finally, any gene clusters whose consensus sequences do not align with any of the databases are characterized as “environmental unknowns”.

After running AGNOSTOS, we identified ∼3.7 million genes overall with an average of about 63 000 genes per sample. Of the 3.7 million predicted genes, ∼2.43 million (66%) were assigned to one of the 574 173 original AGNOSTOS database gene clusters. The remaining genes (∼1.24 million) were clustered separately into 152 634 new gene clusters with more than one gene and 713 940 ‘clusters’ singletons. The new gene clusters (which contained only soil genes) and singletons from the original database that recruited new genes when the soil samples were integrated, were processed through the AGNOSTOS validation, refinement, and classification steps. The new validated set of high-quality gene clusters was integrated with the original clusters to form a new database of ∼598 000 gene clusters containing ∼126 million genes (including ∼2.61 million soil genes). We generated a gene-cluster-by-sample table of gene cluster abundances by using the average coverage depth of each cluster in each sample. As with the taxon-by-sample table (see above), we combined duplicate samples at this point.

### Statistical Analyses

To determine the soil or site characteristics that correlated with the observed variation in the proportional abundances of gene clusters of unknown function in our samples we ran Pearson correlations between the percentage of gene clusters of unknown function in each sample (defined as the number of gene clusters assigned to “genomic unknown” and “environmental unknown” divided by the total number of non-discarded gene clusters), and each of the ten environmental variables we had selected for analysis. We used the same methodology to determine which taxonomic groups (phylum level) were correlated with the proportion of gene clusters of unknown function, after filtering out low-abundance phyla (relative abundance < 0.1 %) to reduce the chance of spurious correlations. For these correlations, we used the false discovery rate method (47) to correct the p-values for multiple tests.

We also identified “dominant” gene clusters of unknown function in soils. A gene cluster was considered dominant based on its abundance and ubiquity across samples, following Delgado-Baquerizo et al. (13). To identify gene clusters of unknown function that were abundant and ubiquitous in soils, we filtered the 550 000 gene clusters in our 48 samples to remove those that were found in fewer than five samples (< 10% of soil samples). This reduced the list of gene clusters to ∼64 000 clusters. Using the Bowtie2 mapping results, we first normalized the per-base coverage by the length of the gene and used this normalized coverage as a proxy for abundance. Then we calculated the median normalized coverage for each gene cluster across all samples. Gene clusters with a median normalized coverage across all samples >0 were considered abundant. After filtering for dominant gene clusters, we identified a total of 43 genomic unknowns and 2461 knowns. We recognize that deeper sequencing would likely have increased the number of dominant gene clusters identified here. However, our goal was not to identify all genes (whether “known” or “unknown”) in each soil, as that is arguably an impossible task given the complexity of soil metagenomes. Rather, our goal was to identify the more abundant and ubiquitous genes found across the set of 48 soils included here.

Because all of the dominant unknowns were classified as genomic unknowns, we wanted to understand which taxa correlated with abundances of these unknowns. To determine this, we ran Spearman correlations between the relative abundances of dominant unknown genes and the abundances of microbial phyla in our soil samples. We used false discovery rate to correct for multiple tests (47). To better understand the functions of the unknowns we ran correlations between each dominant cluster of known function and each dominant unknown cluster using SparCC (48) as implemented in the FastSpar software package (49). To minimize the chance of capturing spurious correlations, we used a p-value cutoff of 0.05, and selected only the strongest correlations (correlation > 0.5) for downstream analysis. Finally, we visualized the results of these gene cluster-to-gene cluster correlations in a co-occurrence network. The network was created in R, using packages R packages tidygraph (https://tidygraph.data-imaginist.com/), and ggraph (https://ggraph.data-imaginist.com/).

### GTDB-classification of dominant gene clusters

To better understand the scope of genomic diversity represented by the dominant unknown gene clusters, we used the Genome Taxonomy Database (GTDB) (50) to assign taxonomy to the contigs containing at least one of each of the 43 dominant unknown gene clusters. We used MMSeqs2 (43) to query sequences against GTDB and assign taxonomy using the MMSeqs2 taxonomic assignment methodology. Briefly, MMSeqs2 uses a computationally efficient method to query each gene on a genome fragment against a reference database and assigns taxonomy to genome fragments containing those genes (i.e., the contigs assigned to one of the 43 dominant unknown genes clusters) based on the 2bLCA taxonomy assignment protocol (51). The 2bLCA method filters hits to a database for quality based on an e-score cut-off (e < 1×10^−12^) and finds the lowest common ancestor of all hits that pass filter. We used GTDB (version 95) as our taxonomy reference database and the following parameters: “--tax-lineage 2 --majority 0.5 --vote-mode 1 --lca-mode 3 --orf-filter 1 --lca-ranks superkingdom,phylum,class,order,family,genus” to run MMSeqs2. From these results, we were able to calculate the number of genomes in GTDB with at least one dominant unknown gene cluster and the most likely taxonomic assignment of each contig based on the taxonomy associated with those genomes in GTDB.

## Results and Discussion

### Abundance of genes of unknown function in soils

We identified ∼550 000 gene clusters (and ∼607 000 singleton clusters for a total of ∼1.16 million clusters) that met the established criteria across the 48 soil samples and calculated the proportion of genes in each sample assigned to each of the four categories (Table 1). On average across the samples, 73.6% of gene clusters were known with Pfam, 10.2% were known without Pfam, 15.8% were genomic unknowns, and 0.4% were environmental unknowns (Figure 1A). The proportion of unknowns varied from 11.9% to 21.7% (Figure 1A). Although not included in downstream analyses as they could not be categorized into the four categories (Table 1), we note that singletons represented a large proportion of identified genes in soils, with nearly twice as many singletons as unknowns (Figure S2A). Singletons also varied from around 13% to 30% of gene clusters per sample, confirming that a large fraction of the genes in soil metagenomics are likely of unknown function (Figure S2A). Importantly, the majority of unknown gene clusters were classified as “genomic unknowns”, suggesting that most unknown gene clusters in these soils represent “low hanging fruit”, that is genes that are unknown (unannotated, hypothetical, or otherwise uncharacterized), but found in genomic databases of sequenced organisms, and therefore more easily characterized than environmental unknowns (i.e., those found exclusively in metagenomes or metagenome-assembled genomes) (11).

**Figure 1:**
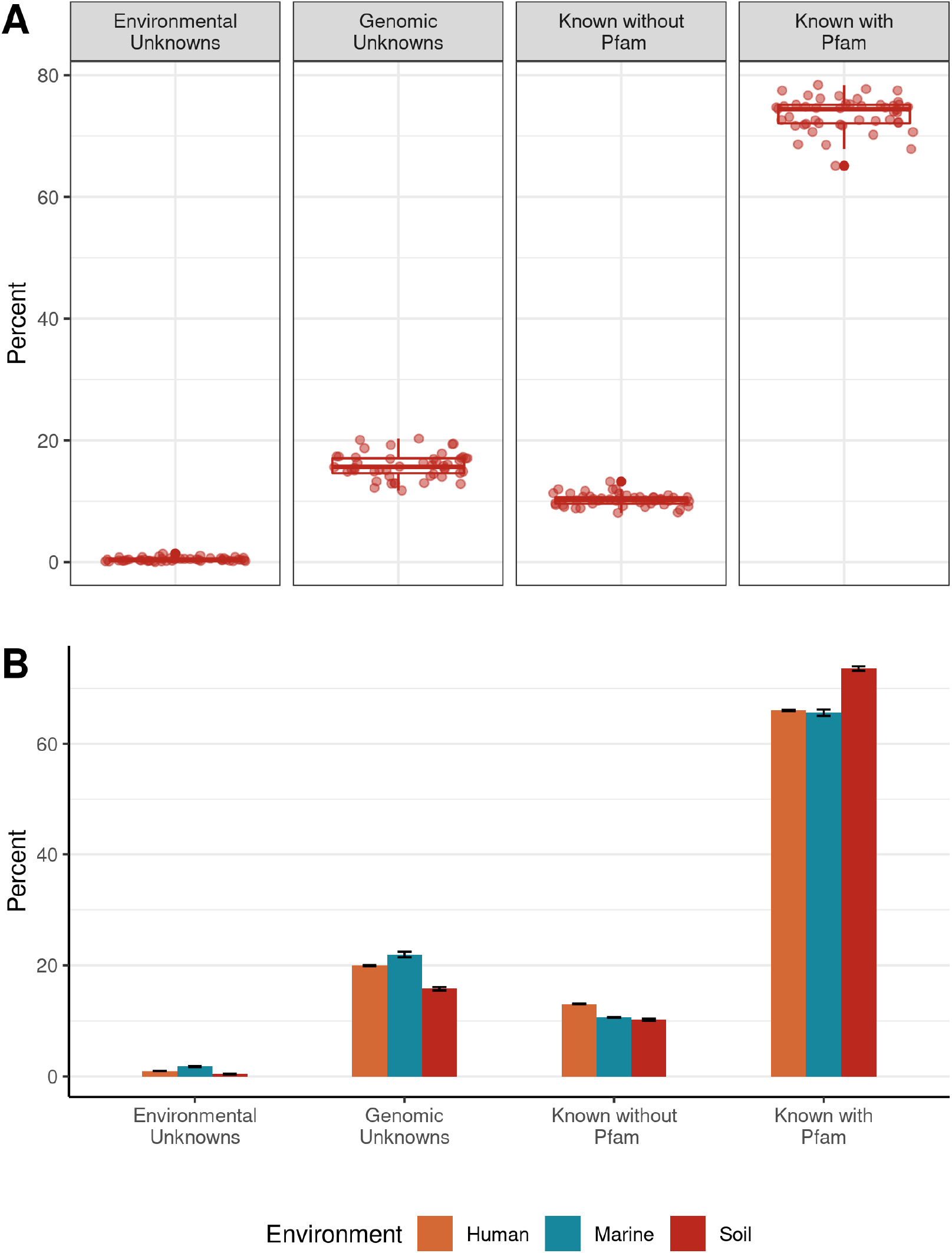
A) The percentage of each gene cluster category in our soil samples. B) A comparison between the percentages of each gene cluster category across metagenomes from three environments: soil, marine, and human-associated. Error bars represent mean standard error across samples.

While the soil metagenomes were found to have a high proportion of genomic unknowns (approximately 12-20% of non-singleton gene clusters), we note that quantifying the specific proportion of genomic unknowns comes with important caveats. First, our ability to detect genes of unknown function in soils is limited by our ability to detect genes in assembled soil metagenomic data. Therefore, we are likely missing many genes from these soil metagenomes as a result of the well-recognized challenges posed by soil metagenomic analyses, and evidenced by the low number of sequences that mapped to our assembled contigs (Table S1) (14,52). The large proportion of singletons we identified in the samples supports this explanation, as they likely represent rarer, genomic and environmental unknowns that are insufficiently captured in our assemblies. These computational challenges limit our ability to detect genes of unknown function that are less abundant or exhibit attributes such as highly variable regions, that make assembly challenging (52–54). Second, the methods we used to classify genes as having a known function yield estimates that are fundamentally conservative. Although a gene may be identified as known, this does not mean that we know much about its function across the wide breadth of microbial diversity (9). As evolution is inherently an iterative process, many proteins from within the same family may exhibit different functions in different organisms and ecological contexts. For example, ammonia monooxygenase and methane monooxygenase genes are assigned to the same Pfam family (AMO), but are functionally distinct (55). Likewise, the similarities between *nifH* genes (involved in nitrogen fixation) and *frxC* genes (involved in light-independent chlorophyll biosynthesis) place them in the same Pfam family (Fer4_NifH), despite coding for distinct proteins (56). Therefore, our estimate of the proportion of genes of unknown function in soil metagenomes is likely an underestimate.

Despite these caveats, we next sought to understand how the proportion of genes of unknown function in soils compared to other metagenomic datasets that were also processed with AGNOSTOS. We compared the total percentages of gene clusters in each category in two additional environments: marine waters and the human microbiome (see refs. (57–61) for assembly statistics). We compared the total number of gene clusters in each category identified in our soil samples to the total number of gene clusters in each category identified in the four marine and two human microbiome metagenomic surveys used in Vanni et al. (11) (Figure 1B). We found that the percentages of gene clusters in each category were consistent across the human microbiome, marine, and soil environments in three of the four gene-cluster categories. On average, 66-74% of the gene clusters in each environment were known with a Pfam designation, 10-13% were known without a Pfam designation, and 16-22% were genomic unknowns. The proportion of environmental unknowns in soils was lower than in marine and human-associated environments, with 0.93% and 1.6% of gene clusters assigned to this category in human and marine environments, respectively, and only 0.32% of soil gene clusters found as environmental unknowns. However, given the diversity of soil metagenomes and the aforementioned challenges associated with assembling genes from soil metagenomes (52,53), this comparatively small proportion of environmental unknowns is likely to be a product of under-sampling the less-abundant genes in soil metagenomes. In support of this explanation, we note the high proportion of singletons in soils (Figure S2B) which are likely composed of a majority of real genes that are rare rather than sequencing artifacts or assembly errors (37).

With the exception of environmental unknowns, the proportions of each gene cluster category were similar across environments. This observation runs counter to our hypothesis that soils would have a higher proportion of unknown genes than other environments. Although it is likely that we are missing many environmental unknowns due to insufficient sequencing and fragmented assemblies, the consistency of our results across environments is supported by their similarity to proportions of each gene category found by Vanni et al. (11) within the GTDB which contains genomes and high quality MAGs from a wide variety of environments. These general similarities in the proportions of genes across environments and genomes suggest that the proportions of unknown and known genes is fairly consistent even across distinct habitat types.

Interestingly, although the proportions of unknown genes in soils were similar to those in marine and human microbiome surveys, this result was not necessarily a product of the same gene clusters being found across all three environments. Of the gene clusters found in the soil samples, ∼ 377 000 (68%) of the clusters were not found in either the human or the marine data sets. Those soil-specific gene clusters were slightly enriched in genomic unknowns (29 %) and environmental unknowns (1.1%) when compared to the set of gene clusters that included human and marine clusters (22.8 % and 0.01 %). These results suggest that although proportions of unknown genes are similar across environments, the soil-specific gene clusters may represent a source of unknown genetic potential that is distinct from that found in either marine or human-associated environments.

### Environmental and microbial groups and genes of unknown functions

Although the majority of gene clusters from assembled contigs in our samples were known, the proportion of unknown non-singleton gene clusters in each soil sample varied from 12-20%. To better understand which environmental factors explained variation in the percentage of unknown genes in our samples, we ran correlations between ten environmental factors (Figure S1C) and the proportion of gene clusters of unknown function in our samples. Of the ten environmental factors, only soil pH (cor = 0.42, p = 0.003) and texture (cor = 0.49, p = 0.001) were significantly correlated with the proportion of gene clusters of unknown function (Figure S3). Importantly, these results were unlikely to be a product of differences in sequencing depth across our samples as the proportion of unknown genes in our samples was not strongly correlated with either sequencing depth (Pearson correlation = 0.26, p = 0.08), or with the total number of genes per sample (Pearson correlation = 0.32, p = 0.03).

Unknown gene clusters were enriched in high-pH soils and finer-textured soils. It is notable that of the ten environmental factors tested, only pH and texture were found to be significant. The most likely explanation for this pattern is that higher pH and finer-textured soils are more likely to harbor a greater fraction of microbes with high proportions of genes of unknown function. This explanation is supported by the analysis of correlations between individual microbial phyla and the proportion of unknown genes in the metagenomes (Figure S4). We identified five phyla that were significantly positively correlated with the percentage of unknowns, and four phyla that were negatively correlated with unknowns. *Crenarchaeota, Gemmatimonadota, Nitrospirota, Entotheonellaeota*, and *Methylomirabilota* were all positively correlated with the proportion of unknown genes. *Bdellovibrionota*, WPS-2 (Eremiobacterota), *Proteobacteri*a, and *Patescibacteria* were all negatively correlated with the percentage of unknowns (Figure S4). Many of the positively correlated phyla are also positively correlated with pH or texture in our dataset (*Gemmatimonadota*: pH – rho = 0.66, *Crenarchaeota*: pH – rho = 0.60, *Methylomirabilota*: texture – rho = 0.47, *Nitrospirota*: pH – rho = 0.47, *Entotheonellaeota*: pH – rho = 0.46, p < 0.05, Spearman correlations) and some of the negatively correlated phyla are negatively correlated with soil pH (WPS-2/Eremiobacterota: pH – rho = -0.55, *Proteobacteria*: pH – rho = -0.50, p < 0.05, Spearman correlations). While correlations do not imply causal relationships, the most parsimonious explanation for these results is that higher pH and finer textured soils are more likely to harbor certain phyla that have higher proportions of genes of unknown function and serve as ripe targets for explorations of novel genetic diversity.

### A “hit list” of ubiquitous and abundant genes of unknown function in soils

To identify the dominant gene clusters found in our samples, we filtered our data to select only the most abundant and ubiquitous gene clusters in soil samples (see Methods). Applying these criteria yielded 43 gene clusters of unknown function, all of which fell into the genomic unknown category, and 2461 gene clusters of known function (Figure 2). For simplicity, we hereafter refer to the 43 abundant and ubiquitous gene clusters as “dominant” unknown gene clusters.

**Figure 2:**
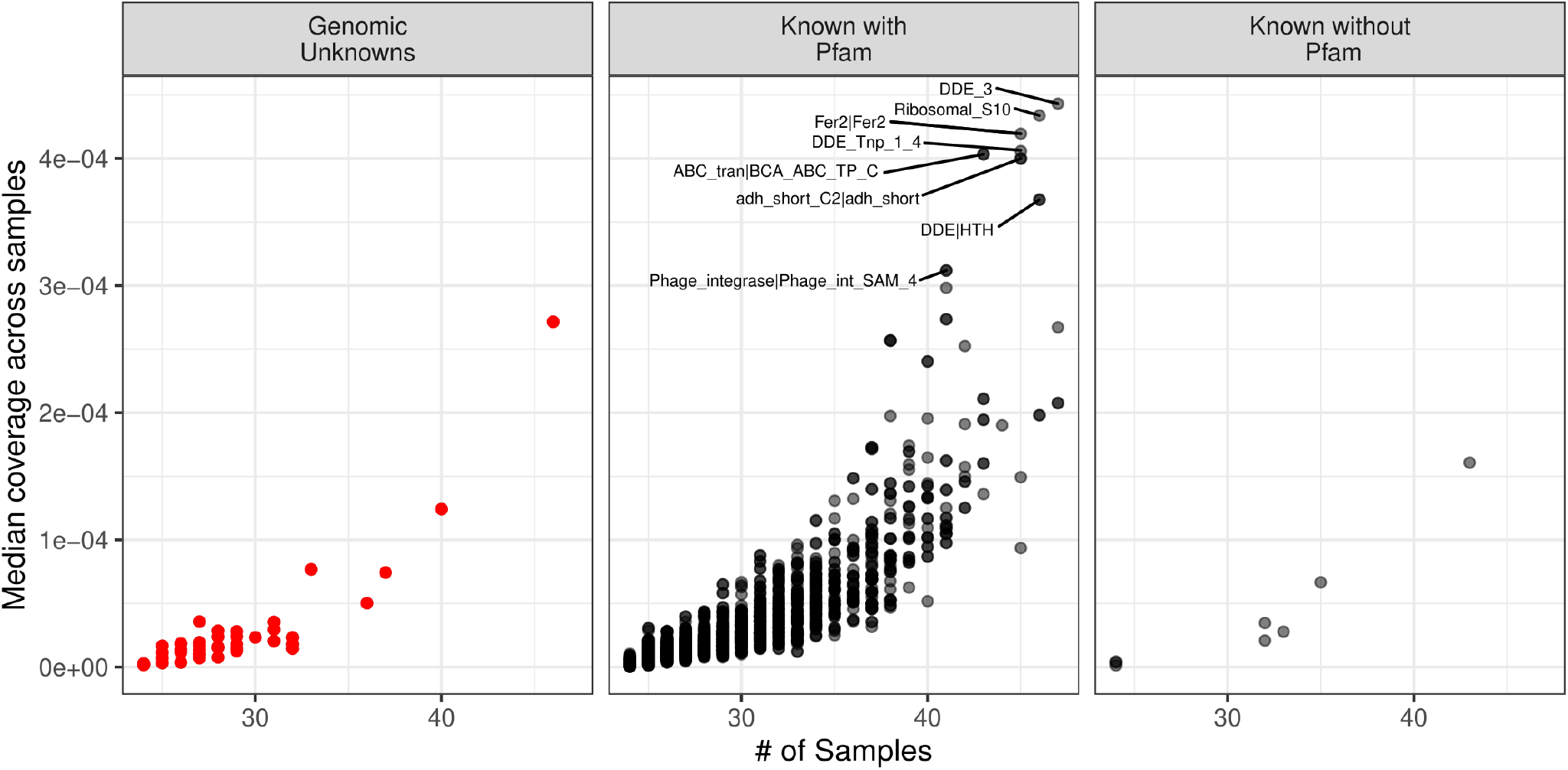
The “dominant” gene clusters in our samples. Each point represents one gene cluster. The x-axis shows the number of soil samples in which each gene cluster was found (“ubiquity”) and the y-axis shows the median coverage of each gene cluster across those samples (“abundance”). The most dominant clusters are in the top right corner of each plot, while less dominant clusters are in the bottom left corner of each plot. We identified 43 dominant unknowns in our samples (red) and 2461 knowns (black). Environmental unknowns are omitted from this figure, as no environmental unknowns were sufficiently abundant (mean coverage > 0) and ubiquitous (present in > 10% of samples) to be considered a “dominant” gene cluster. Pfam designations for some of the most abundant and ubiquitous known genes are indicated in the “Known with Pfam” panel.

To better understand the genomic and ecological context of the dominant unknown gene clusters, we ran correlations between the clusters and phyla in our samples (Figure 3). Positive correlations between the dominant gene clusters and taxa ranged from 0.41-0.77. While some of the gene clusters were positively correlated with many taxa (e.g., GC34767203 correlated with 8 phyla), others were only significantly correlated with one or two phyla (e.g., GC17275105). The phyla that had the largest numbers of positive correlations to the dominant unknown gene clusters included the Unclassified Phylum RCP2-54, *Verrucomicrobiota, Proteobacteria, Acidobacteriota*, and *Planctomycetota* (Figure 3). Many of the phyla that were positively correlated with dominant unknown gene clusters were lineages that are typically found in high abundances in soils (13). While many of these lineages, such *Acidobacteria* or *Verrucomicrobia*, contain large numbers of uncultivated taxa (62) some of these phyla come from lineages that have been reasonably well-studied (e.g. *Proteobacteria*). While positive correlations do not necessarily imply the presence of these genes in genomes from members of these phyla, these well-studied taxa potentially represent a straightforward and simple route to discovering the functions of these gene clusters as many of the members of these phyla are reasonably well-represented in culture collections and genomic databases. By searching for these dominant unknowns in genome databases, and cross-referencing the results with strains that are available in culture collections, it may be feasible characterize the functions of dominant unknown genes in soils using targeted experiments, as demonstrated by Price et al. (9).

**Figure 3:**
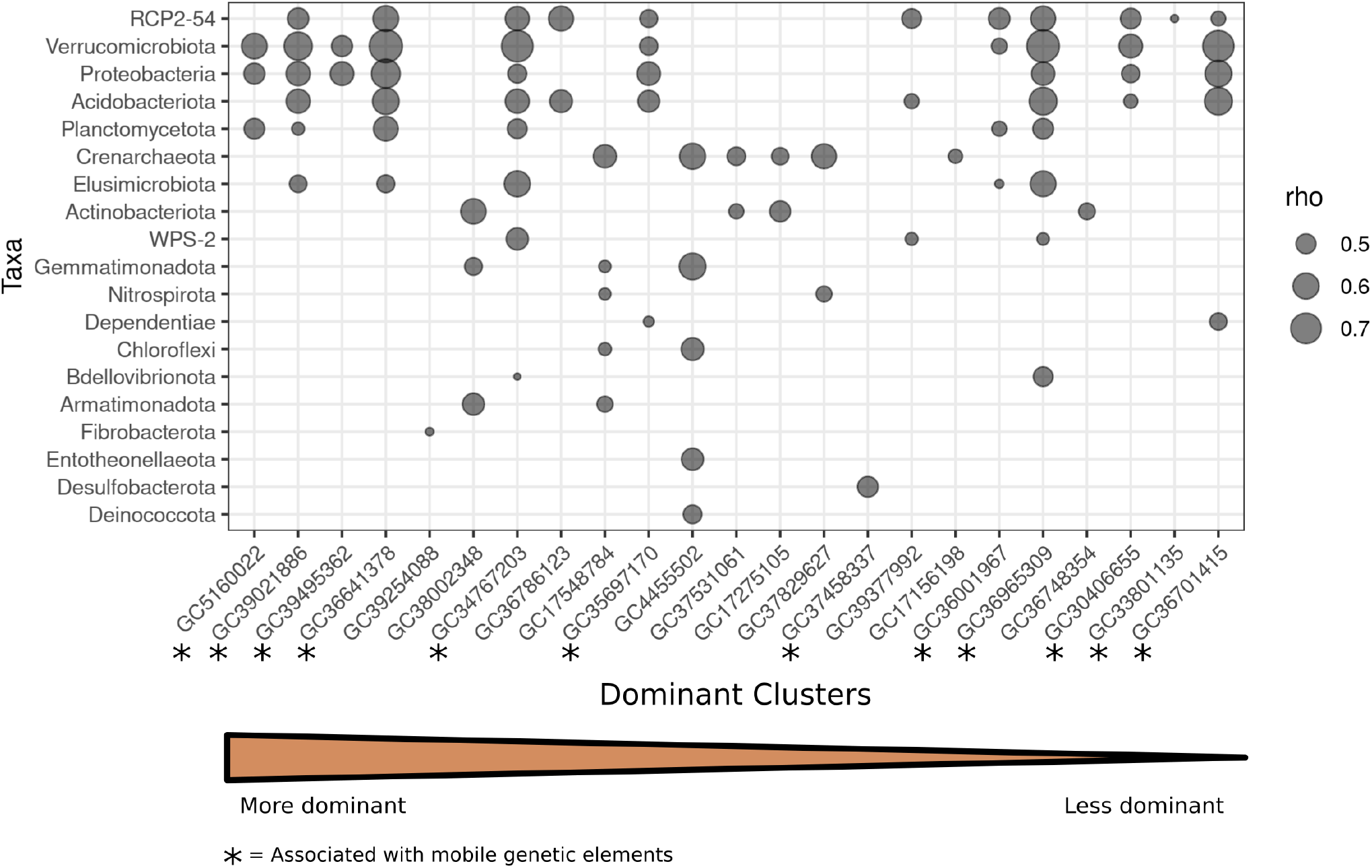
Positive correlations between dominant unknown gene clusters and specific bacterial and archaeal phyla across the 48 soil samples included in this study. Each point represents a significant Spearman correlation (p < 0.05) between one of the 43 unknown clusters (x-axis) and a microbial phylum (y-axis). Phyla are arranged in order of increasing number of significant correlations to dominant clusters (bottom to top). Clusters are arranged in decreasing order of dominance (left to right). The size of each point represents the strength of the rho value. Gene clusters that were also found to be strongly associated with many mobile genetic elements (see Figure 5) are indicated with asterisks.

To further understand the genomic context of the dominant unknown gene clusters, we searched for the contigs associated with our 43 dominant unknown gene clusters in the GTDB (50) and used a consensus taxonomy assignment approach to assign taxonomy to each contig. Our goal was to try to identify which taxa might harbor these dominant unknown genes by searching for these genes in a curated genome database, versus simply assessing correlations between taxa and cluster abundances directly from shotgun data (Figure 3). The median number of genomes with matches to each dominant unknown gene cluster was 175, ranging from 38 (GC36965309) to 796 (GC38645763) genomes per dominant cluster and, in general, clusters that were unique to our soil data set (i.e., not found in either the marine or human microbiome datasets) had fewer hits to genomes in the database (Figure 4). Confirming the correlation-based results described above and in Figure 3, the most well-represented phyla across the dominant unknown gene clusters were those that are typically abundant in soils (e.g., *Proteobacteria, Acidobacteria*, and *Actinobacteria*) (Figure 4). However, there are some important discrepancies between the taxonomic results from the GTDB-database and the correlation results (Figure 3). Many of the less well-studied lineages that were correlated with the abundances of particular phyla in our soil dataset (Figure 3) are rare or absent from the GTDB analysis (e.g., Unclassified Phylum RCP2-54 and *Verrucomicrobia*). This is likely due to the fact that some of these lineages are not well represented in GTDB. For example, there are no sequenced genomes or MAGs for RCP2-52 and, even other phyla for which far more genomes are available in GTDB, genomes from representative soil lineages may not be well-represented in GTDB. For example, the phylum Actinobacteria, one of the best-represented lineages in GTDB and one of the most abundant phyla in soils globally (13) has 24 602 entries in GTDB, but only 427 (1.7%) are representatives isolated from soil. Thus, we expect that some of these dominant unknown genes in soil may be associated with lineages that are reasonably common in soil, but under-represented in GTDB and comparable genomic databases. Alternatively, the observed discrepancies may be the result of spurious correlations with the correlation-based analyses (Figure 3). However, the overall consistency between the two results suggests that the majority of these relationships are likely robust, and that a paired approach using multiple methods of taxonomic assignment can be a productive method to generate hypotheses about the taxa associated with genes of unknown function.

**Figure 4:**
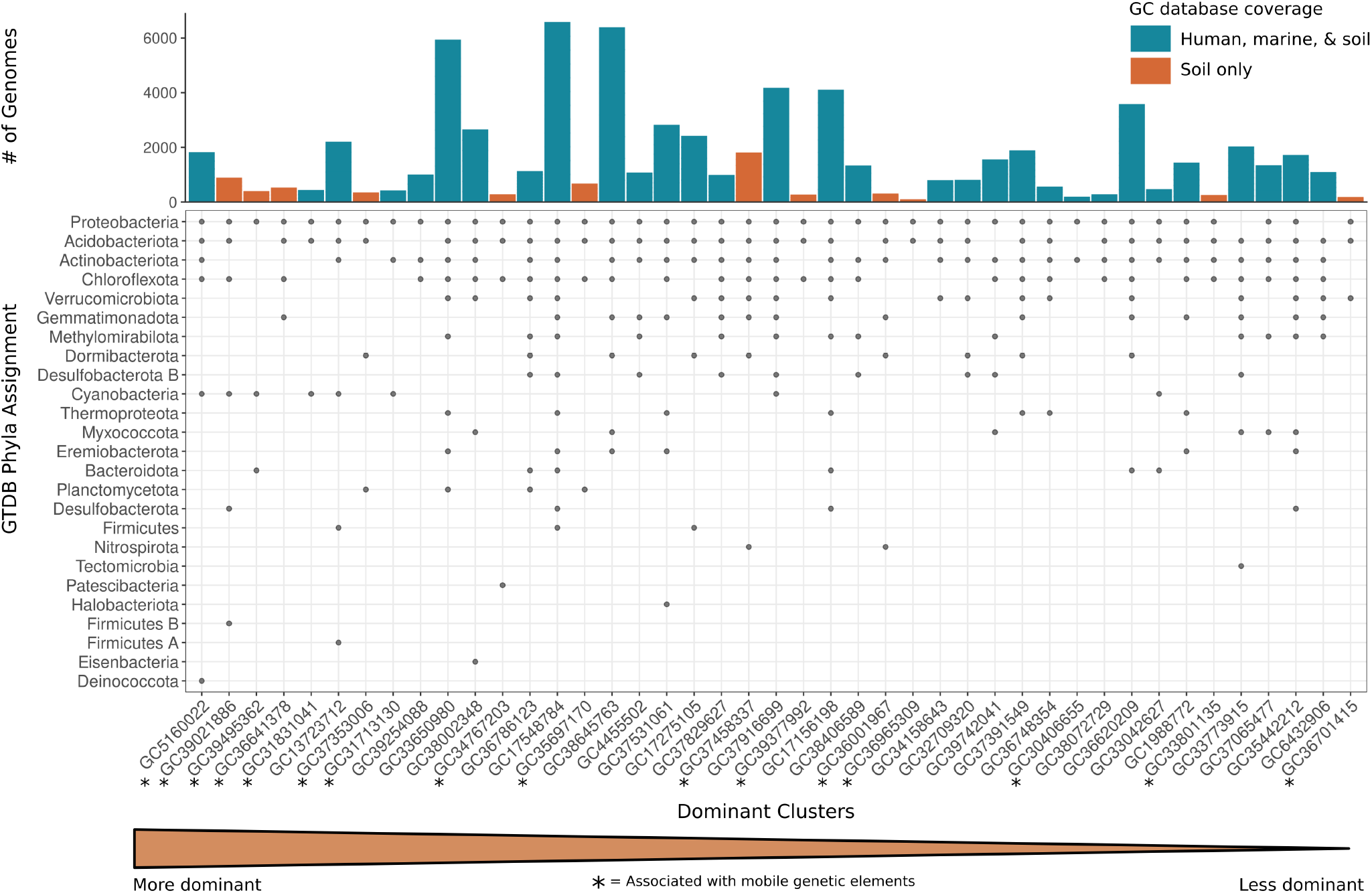
The taxonomic representation (phylum level) of contigs in dominant unknown gene clusters mapped against the GTDB genome database. Contigs within each dominant unknown gene cluster were assigned to a particular phylum based on best-hit consensus taxonomy assignments of the genes within that contig using the 2blca method as implemented in mmseqs2. In the lower panel, a point indicates the presence of at least one contig from the gene cluster (x-axis) being assigned to a particular phylum (y-axis). In upper panel, bars indicate the number of genomes that contigs within a given gene cluster mapped to (a hit to a genome was counted only for hits passing the filter threshold e-score < 1×10^−12^). The colors of the bars indicate whether or not a particular gene cluster was also found in marine or human microbiome datasets (blue) or if the cluster was only found in soils (orange). Dominant clusters that were also found to be strongly associated with many mobile genetic elements (see Figure 5) are indicated with asterisks.

In addition to determining the potential taxonomic affiliation of the dominant genes of unknown function, we also sought to determine their potential functions. We did so by assessing relationships between known (annotated) gene clusters and the dominant “unknown” gene clusters across the 48 metagenomes, following the working assumption that annotated and unknown genes which are correlated may have related functions (63,64). In total, 16 of the 43 gene clusters had at least one positive correlation with known gene clusters above the established threshold. All 16 of these genomic unknowns were correlated with mobile genetic elements such as transposases, reverse transcriptases, or phage integrases (Table 2). These mobile genetic element-associated genes made up the majority of “known” genes correlated with each of the 16 dominant genes of unknown function with (Table 2). These patterns are evident from a co-occurrence network visualization of the associations between genes of known function and those of unknown function (Figure 5, see Figure S5 for a version of this network with labeled known gene clusters). Importantly, many of the gene clusters associated with mobile genetic elements were also positively correlated with more bacterial and archaeal phyla (Figure 3), and their positive correlations with mobile genetic elements may suggest a mechanism as to why they are so taxonomically widespread.

**Figure 5:**
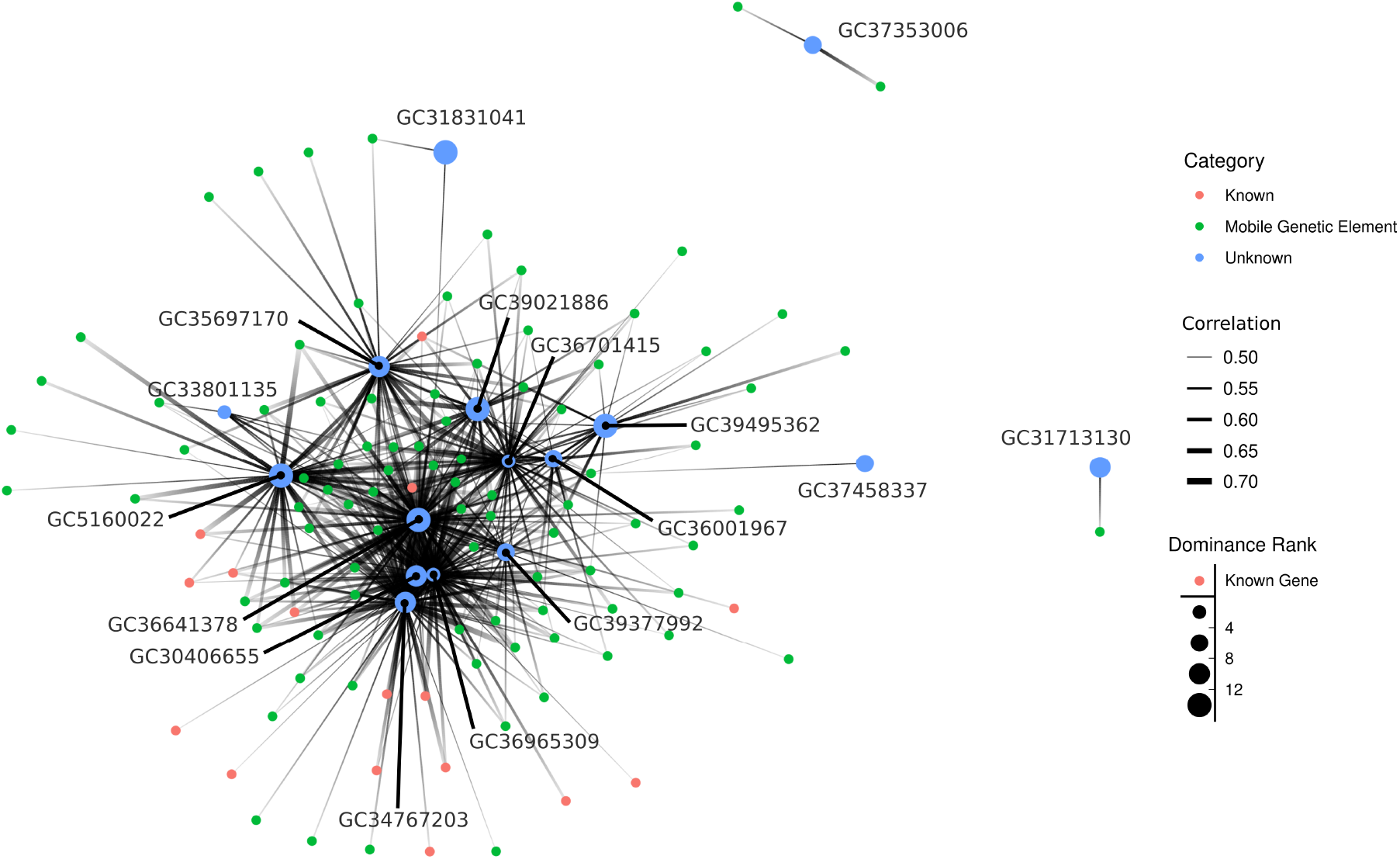
A co-occurrence network showing significant positive Spearman correlations (rho > 0.6) between dominant unknown gene clusters and dominant known gene clusters. Unknown genes are indicated with the blue points, and the size of the points indicates how dominant they are relative to other dominant unknowns, with larger circles indicating a greater level of dominance. Known genes are displayed in green. Green points represent known genes that are mobile elements while all other known genes are displayed in red. The thickness of the edges represents the strength of the rho value. A version of this figure with all points labeled with their Pfam designations can be found in the supplementary material (Figure S5).

Mobile genetic elements represent one of the main categories of unexplored genetic diversity (65) and are thought to be one of the most abundant functional gene categories across the tree of life (66). Mobile elements are associated with elevated genomic plasticity and accelerated biological diversification (66). While traditionally associated with antibiotic resistance genes, one study of integrons across a broad diversity of metagenomes ranging from oceans, to guts, to soils, found that many are likely to be toxin/anti-toxin genes, but that most integron genes are of unknown function (65). Their results are consistent with the findings from this study, which suggest that many of the most dominant genes of unknown function in soils are correlated with the abundances of transposases, recombinases, resolvases, phage integrases, and other mobile genetic elements. Clearly further investigations into the ecology, function and genomic contexts of mobile genetic elements are warranted, considering that many of the dominant unknown genes we identified from soil tend to co-occur with genes coding for mobile genetic elements.

## Conclusions

We have demonstrated that genes of unknown function comprise a substantial portion of the genes in soils and we are likely underestimating their true abundances due to challenges associated with adequately capturing the full extent of soil metagenomic diversity. Most notably, genes of unknown function were more abundant in some soils than others and this variance was, in part, predictable from soil edaphic properties and the taxonomic composition of the soil microbial communities. Finally, we were able to identify the most dominant genes of unknown function in soil, defined as those genes that are abundant and ubiquitous across a wide range of soil types. We find that these dominant genes of unknown function are primarily found in typically abundant soil lineages that are less-well studied such as *Verrucomicrobia* and *Acidobacteria*, with the majority of these dominant genes associated with mobile genetic elements. Our work demonstrates the utility of investigating genes of unknown function and represents a first step towards “ecological annotation” of the dominant genes of unknown function in soils to elucidate what taxa are likely to have these genes, what environments they are most likely to be found in, and the putative functions of the unknown genes that represent an important fraction of soil metagenomes.

## Acknowledgments

We would like to thank Christopher Miller and Sarah Bagby for helpful comments on the manuscript and methods. We would like to acknowledge the contribution of the Biomes of Australian Soil Environments (BASE) consortium in the generation of data used in this publication. The BASE project was supported by funding from Bioplatforms Australia through the Australian Government National Collaborative Research Infrastructure Strategy (NCRIS). We would also like to acknowledge the use of the Atlas of Living Australia, spatial data <u>grid.506668.b</u> We also acknowledge the computer resources at MareNostrum and the technical support provided by Barcelona Supercomputing Center (RES-AECT-2014-2-0085), the BMBF-funded de.NBI Cloud within the German Network for Bioinformatics Infrastructure (de.NBI) (031A537B, 031A533A, 031A538A, 031A533B, 031A535A, 031A537C, 031A534A, 031A532B), the University of Oxford Advanced Research Computing (http://dx.doi.org/10.5281/zenodo.22558). CV was supported by the Max Planck Society.

