## Supplemental Tables and Figures for "An ecological perspective on microbial genes of unknown function in soil"

**Supplemental Tables and Figures for:**
### An ecological perspective on microbial genes of unknown function in soil

Hannah Holland-Moritz*[1], Chiara Vanni[2], Antonio Fernandez-Guerra[3], Andrew Bissett[4], and Noah Fierer[5]

[1] Department of Natural Resources and the Environment, University of New Hampshire, Durham, NH, USA
[2] Microbial Genomics and Bioinformatics Research Group, Max Planck Institute for Marine Microbiology, Celsiusstraße 1, 28359, Bremen, Germany
Jacobs University Bremen, Campus Ring 1, 28759 Bremen, Germany
[3] Lundbeck GeoGenetics Centre, The Globe Institute, University of Copenhagen, Copenhagen, Denmark
[4] CSIRO, Oceans and Atmosphere, Hobart, Tasmania, Australia
[5] Department of Ecology and Evolutionary Biology, Cooperative Inst. for Research in Environmental Sciences, University of Colorado, Boulder, USA

#### Supplemental Tables and Figures
**(Some of these files are larger than a page, please find them in the additional supplementary files provided.)**

| SampleID | Australian Microbiome Initiative ID | Sequencing Depth (millions of reads) | N50 (Kbp) | N75 (Kbp) | L50 (K) | L75 (K) | Assembly Length (Mbp) | % Mapped |
|---|---|---|---|---|---|---|---|---|
| Sample_8268 | 102.100.100/8268 | 154.9 | 0.80 | 0.6 | 180.15 | 349406.0 | 954.8 | 9.165 |
| Sample_8154 | 102.100.100/8154 | 151.7 | 0.70 | 0.6 | 127.35 | 224546.0 | 492.5 | 2.488 |
| Sample_7061 | 102.100.100/7061 | 68.7 | 0.80 | 0.6 | 37.75 | 70844.5 | 185.4 | 3.284 |
| Sample_15977 | 102.100.100/15977 | 49.7 | 0.80 | 0.6 | 41.35 | 81097.5 | 229.0 | 5.635 |
| Sample_12876 | 102.100.100/12876 | 44.8 | 1.30 | 0.8 | 89.70 | 215682.0 | 493.4 | 26.773 |
| Sample_15955 | 102.100.100/15955 | 43.0 | 0.70 | 0.6 | 42.90 | 78370.0 | 93.4 | 1.711 |
| Sample_12824 | 102.100.100/12824 | 39.3 | 0.90 | 0.6 | 69.40 | 140417.0 | 211.8 | 7.442 |
| Sample_12939 | 102.100.100/12939 | 39.0 | 0.90 | 0.6 | 59.80 | 117337.0 | 169.9 | 5.135 |
| Sample_8455 | 102.100.100/8455 | 38.1 | 0.80 | 0.6 | 50.00 | 100352.0 | 144.7 | 4.604 |
| Sample_12432 | 102.100.100/12432 | 37.9 | 0.80 | 0.6 | 46.80 | 86171.0 | 104.6 | 2.353 |
| Sample_7861 | 102.100.100/7861 | 36.9 | 0.80 | 0.6 | 19.00 | 36835.5 | 99.2 | 3.115 |
| Sample_7912 | 102.100.100/7912 | 36.1 | 0.80 | 0.6 | 60.70 | 112154.0 | 143.4 | 3.955 |
| Sample_12618 | 102.100.100/12618 | 35.5 | 0.80 | 0.6 | 21.00 | 38796.0 | 47.3 | 0.946 |
| Sample_7069 | 102.100.100/7069 | 34.3 | 0.80 | 0.6 | 64.30 | 117240.0 | 142.4 | 3.459 |
| Sample_13276 | 102.100.100/13276 | 34.3 | 0.70 | 0.6 | 9.65 | 17093.0 | 37.1 | 0.556 |
| Sample_15885 | 102.100.100/15885 | 34.3 | 1.00 | 0.7 | 23.45 | 49839.0 | 170.2 | 7.166 |
| Sample_7888 | 102.100.100/7888 | 33.9 | 0.90 | 0.6 | 51.30 | 108585.0 | 174.0 | 7.608 |
| Sample_12830 | 102.100.100/12830 | 33.6 | 0.90 | 0.6 | 64.10 | 123354.0 | 172.6 | 7.596 |
| Sample_8128 | 102.100.100/8128 | 32.4 | 0.70 | 0.6 | 31.30 | 56834.0 | 66.9 | 1.585 |
| Sample_12495 | 102.100.100/12495 | 32.3 | 0.70 | 0.6 | 12.65 | 22842.0 | 53.0 | 1.193 |
| Sample_9462 | 102.100.100/9462 | 32.2 | 0.90 | 0.6 | 41.80 | 81305.0 | 117.2 | 4.788 |
| Sample_7892 | 102.100.100/7892 | 31.9 | 0.90 | 0.6 | 70.80 | 137889.0 | 199.7 | 8.242 |
| Sample_19457 | 102.100.100/19457 | 31.0 | 0.70 | 0.6 | 7.85 | 13841.5 | 30.0 | 0.430 |
| Sample_7063 | 102.100.100/7063 | 30.2 | 0.70 | 0.6 | 27.60 | 47882.0 | 50.9 | 0.912 |
| Sample_7902 | 102.100.100/7902 | 30.2 | 1.10 | 0.7 | 36.90 | 88631.0 | 173.3 | 13.051 |
| Sample_14183 | 102.100.100/14183 | 29.9 | 0.60 | 0.6 | 5.30 | 9028.0 | 17.7 | 0.189 |
| Sample_12485 | 102.100.100/12485 | 29.4 | 0.80 | 0.6 | 29.90 | 55526.0 | 68.1 | 2.938 |
| Sample_12481 | 102.100.100/12481 | 29.3 | 0.70 | 0.6 | 10.00 | 17851.0 | 40.0 | 0.886 |
| Sample_19517 | 102.100.100/19517 | 29.1 | 0.80 | 0.6 | 17.55 | 32803.5 | 84.6 | 2.616 |
| Sample_8088 | 102.100.100/8088 | 28.8 | 1.10 | 0.7 | 46.00 | 101274.0 | 193.5 | 13.386 |
| Sample_19453 | 102.100.100/19453 | 27.8 | 0.70 | 0.6 | 7.55 | 13220.0 | 28.0 | 0.522 |
| Sample_7829 | 102.100.100/7829 | 27.7 | 0.75 | 0.6 | 19.05 | 34330.0 | 81.1 | 2.341 |
| Sample_12838 | 102.100.100/12838 | 27.6 | 0.80 | 0.6 | 21.60 | 39969.0 | 49.4 | 1.628 |
| Sample_8130 | 102.100.100/8130 | 27.1 | 0.70 | 0.6 | 26.00 | 47343.0 | 56.7 | 1.938 |
| Sample_8164 | 102.100.100/8164 | 26.0 | 0.80 | 0.6 | 34.50 | 63163.0 | 78.8 | 3.117 |
| Sample_12523 | 102.100.100/12523 | 25.3 | 0.80 | 0.6 | 12.60 | 23216.0 | 58.5 | 2.016 |
| Sample_19501 | 102.100.100/19501 | 24.6 | 0.70 | 0.6 | 9.20 | 16329.5 | 35.9 | 0.941 |
| Sample_13729 | 102.100.100/13729 | 24.0 | 0.90 | 0.7 | 34.00 | 71654.0 | 115.9 | 8.102 |
| Sample_8142 | 102.100.100/8142 | 23.9 | 0.70 | 0.6 | 7.35 | 12828.5 | 26.8 | 0.505 |
| Sample_12560 | 102.100.100/12560 | 23.6 | 0.90 | 0.7 | 40.80 | 83093.0 | 130.8 | 10.219 |
| Sample_12497 | 102.100.100/12497 | 20.4 | 0.70 | 0.6 | 4.65 | 8055.0 | 16.8 | 0.342 |
| Sample_12467 | 102.100.100/12467 | 17.3 | 0.70 | 0.6 | 4.40 | 7720.0 | 16.7 | 0.562 |
| Sample_7075 | 102.100.100/7075 | 16.0 | 0.70 | 0.6 | 3.35 | 5762.0 | 11.6 | 0.366 |
| Sample_9466 | 102.100.100/9466 | 15.2 | 0.65 | 0.6 | 2.15 | 3693.0 | 7.4 | 0.172 |
| Sample_12465 | 102.100.100/12465 | 14.0 | 0.65 | 0.6 | 1.95 | 3458.5 | 7.0 | 0.255 |
| Sample_12469 | 102.100.100/12469 | 13.5 | 0.70 | 0.6 | 3.05 | 5247.5 | 10.7 | 0.340 |
| Sample_8497 | 102.100.100/8497 | 13.4 | 0.60 | 0.6 | 2.15 | 3733.5 | 7.5 | 0.260 |
| Sample_9450 | 102.100.100/9450 | 11.8 | 0.70 | 0.6 | 4.00 | 7098.5 | 15.8 | 0.900 |

**Table S1:** Table showing sample accession IDs, sequencing depth and assembly statistics for each sample.

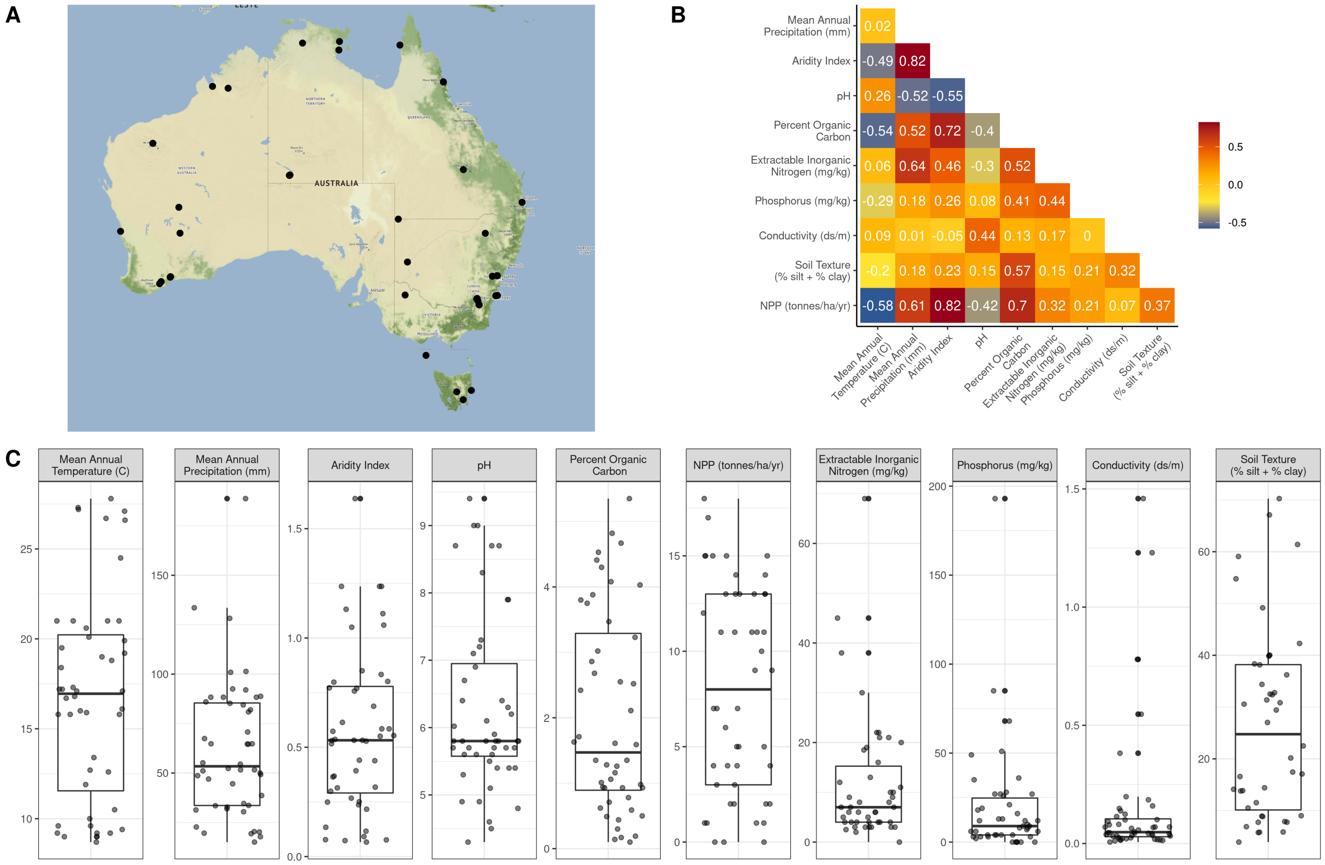

**Figure S1:** A summary of the samples used in this dataset and their contextual metadata. A) a map of Australia showing the locations of the 48 samples. B) Correlations among the 10 environmental factors we selected for downstream analysis. C) The ecological gradients spanned by each of the samples for each metadata category.

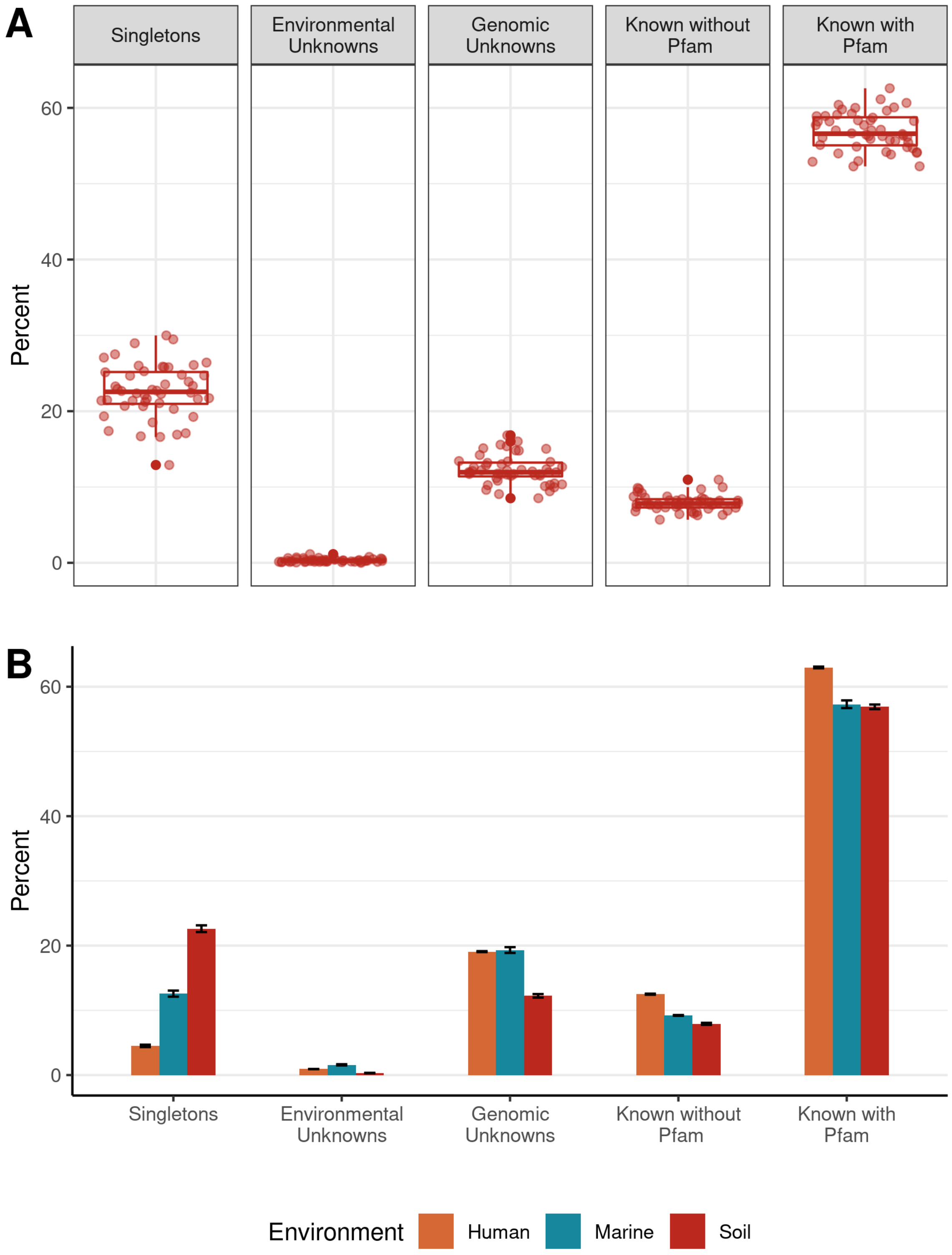


**Figure S2:** A) The percentage of each gene cluster category in our soil samples as well as the percentages of singleton clusters. B) A comparison between the percentages of each gene cluster category across metagenomes from three environments, soil, marine, and human-associated. Error bars represent mean standard error across samples.

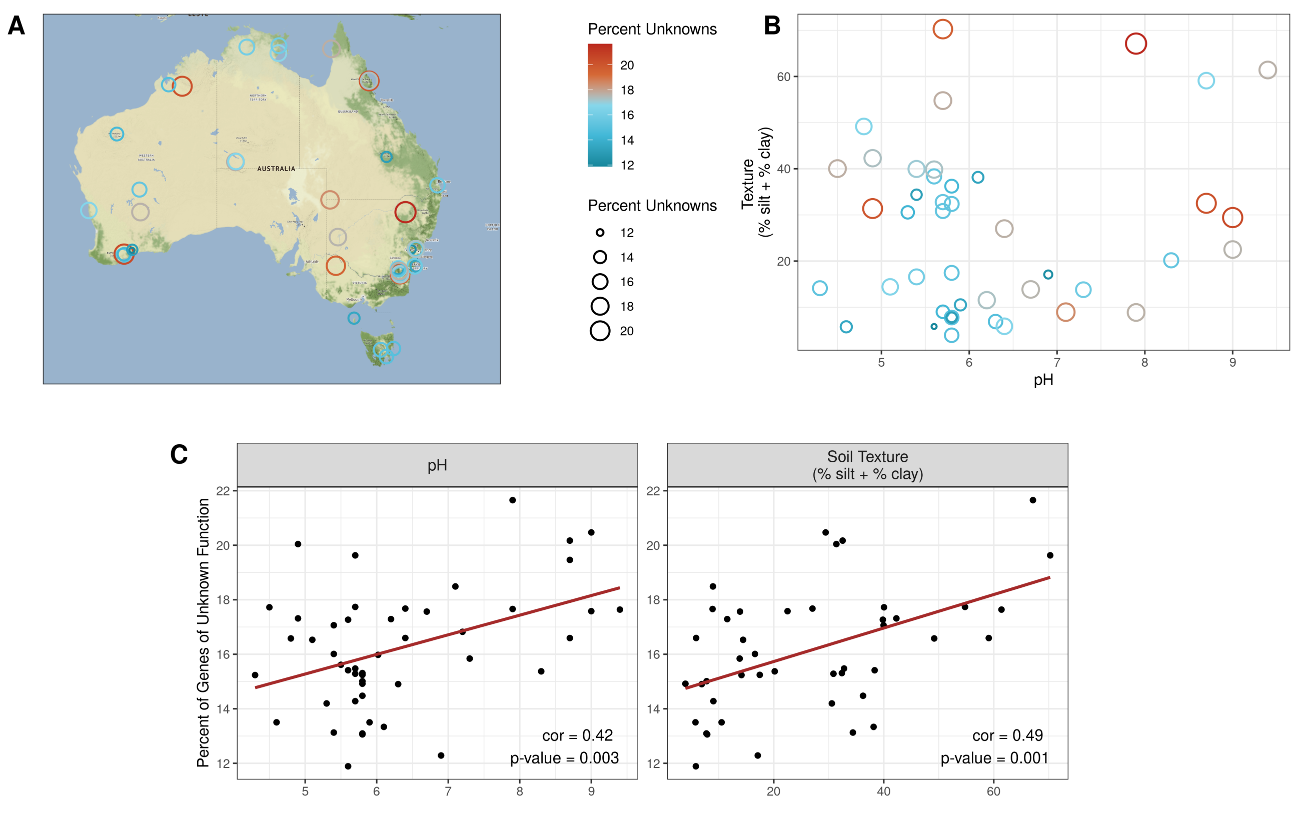


**Figure S3:** A) A map showing the locations of the 48 soil samples with the color and size of the points indicating the proportion of unknowns in each sample. Larger, red points indicate a higher percentage of unknown gene clusters, small, blue points represent a lower percentage of unknown gene clusters. B) The relationship between soil texture and pH in our data. Point color and size corresponds to that in panel A. C) Significant Pearson correlations between the percent of gene clusters of unknown function and pH and soil texture (% silt + % clay).

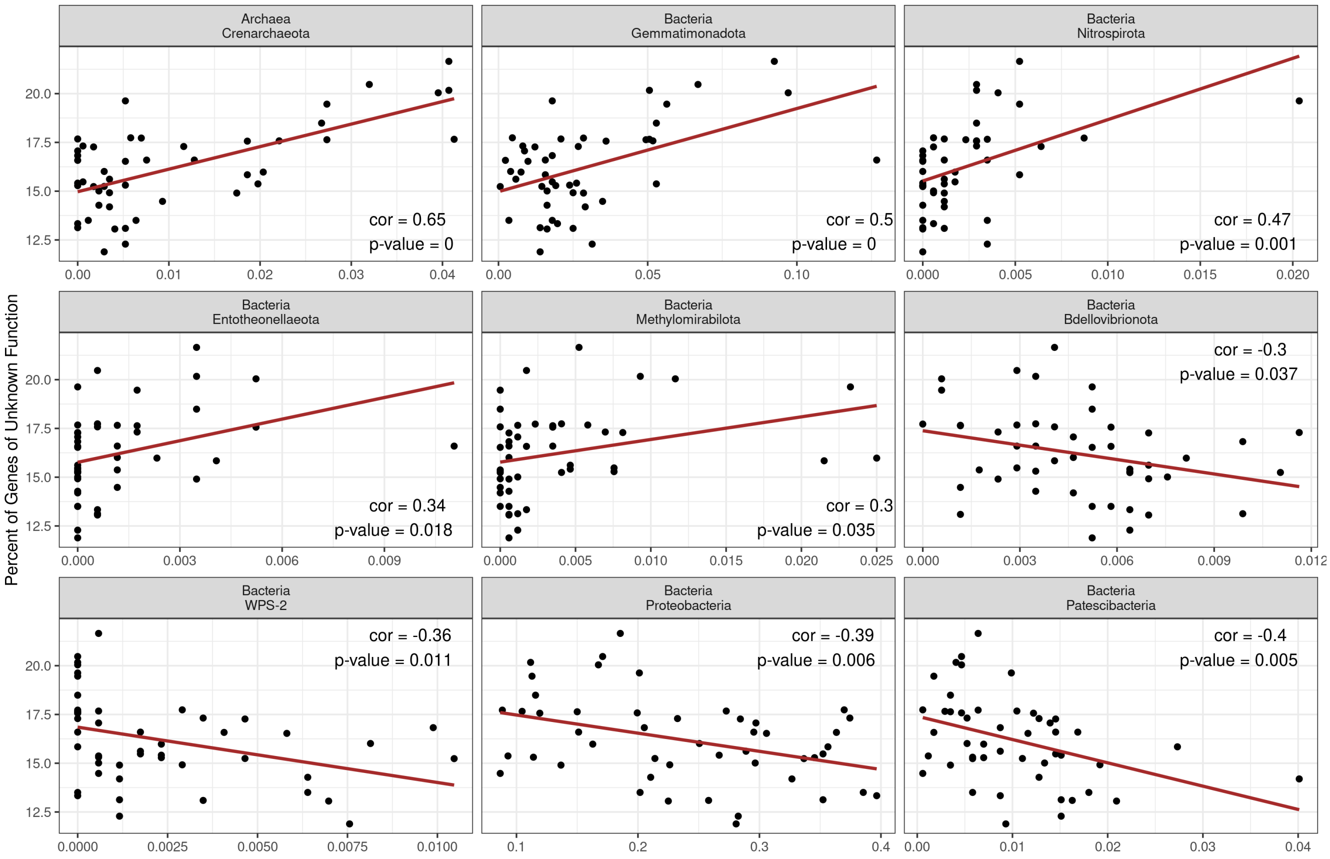


**Figure S4:** Significant correlations between the relative abundances of microbial phyla in the soil samples and the percent of genes of unknown function. The relative abundance of each phyla (% of reads) is represented on the x-axis, while the percent of genes of unknown function in each sample is represented by the y-axis.

**Figure S5:** A more detailed version of Figure 5 with labels for all genes with functional annotations. Labels give the assigned Pfam designation for each gene cluster.

**Table S2:** Table showing which dominant known gene clusters were significantly and strongly (> 0.5) correlated with the dominant unknown gene clusters and their Pfam annotation. Genes that are or are related to mobile genetic elements are indicated in the "General Function" column. We categorized genes semi-manually as mobile genetic elements by searching for key terms. A gene was considered related to mobile genetic element if its Pfam name contained one of the following search terms: "Transposase" - a known transposase, "Tnp" – a common abbreviation for transposases in Pfam, "Phage_int" – phage integrases, "rve" – retroviral integrases, "Recombinase" – a known recombinase, "Resolvase" – a known resolvase, "RVT" – reverse transcriptases, "GIIM" - Group II intron, maturase-specific domain, involved in reverse transcription and splicing, "IstB" – family of insertion sequences, thought to be related to transposases.
