## Supplementary material for "An ecological perspective on microbial genes of unknown function in soil": Table S2

| Gene Clusters | Associated Pfam Cateogories | General Function |
|---|---|---|
| **GC5160022** | | |
| GC15813830 | DEDD_Tnp_IS110\|Transposase_20 | Mobile genetic elements |
| GC18411621 | DDE_Tnp_1_6\|DUF772 | Mobile genetic elements |
| GC19465353 | DDE_Tnp_1_2 | Mobile genetic elements |
| GC22200452 | DDE_Tnp_1\|DUF772 | Mobile genetic elements |
| GC23019517 | DDE_Tnp_1_4 | Mobile genetic elements |
| GC23227492 | GIIM\|RVT_1 | Mobile genetic elements |
| GC24192253 | HTH_32 | |
| GC30561759 | Phage_integrase\|Phage_int_SAM_4 | Mobile genetic elements |
| GC35526541 | HTH\|HTH | |
| GC38440389 | Y2_Tnp\|Zn_Tnp_IS91 | Mobile genetic elements |
| GC38580544 | Transposase_20 | Mobile genetic elements |
| GC465251 | ACR_tran\|ASL_C2 | |
| GC5059750 | Y2_Tnp\|Zn_Tnp_IS91 | Mobile genetic elements |
| GC8673204 | ACR_tran | |
| GC11494356 | GIIM\|RVT_1 | Mobile genetic elements |
| GC13014417 | Recombinase\|Resolvase | Mobile genetic elements |
| GC23310339 | DDE_3 | Mobile genetic elements |
| GC24933148 | DDE_Tnp_IS1595\|Zn_Tnp_IS1595 | Mobile genetic elements |
| GC25354868 | HTH_32\|HTH_33 | |
| GC27144963 | Transposase_20 | Mobile genetic elements |
| GC31612606 | GIIM\|RVT_1 | Mobile genetic elements |
| GC31969864 | DDE_Tnp_1\|DUF772 | Mobile genetic elements |
| GC31986313 | DEDD_Tnp_IS110\|Transposase_20 | Mobile genetic elements |
| GC33500111 | DEDD_Tnp_IS110\|Transposase | Mobile genetic elements |
| GC36248678 | GIIM\|RVT | Mobile genetic elements |
| GC36543908 | DDE_5 | Mobile genetic elements |
| GC37723193 | DDE_3 | Mobile genetic elements |
| GC39015032 | DEDD_Tnp_IS110\|Transposase | Mobile genetic elements |
| GC39316024 | GIIM\|RVT | Mobile genetic elements |
| GC39642572 | GIIM\|RVT | Mobile genetic elements |
| GC39817105 | Phage_integrase | Mobile genetic elements |
| GC39878062 | ACR_tran | |
| GC5906625 | DDE_3\|HTH_32 | Mobile genetic elements |
| GC26926321 | DDE_Tnp_1_2 | Mobile genetic elements |
| GC27189918 | Y2_Tnp | Mobile genetic elements |
| GC35916463 | TnpB_IS66 | Mobile genetic elements |
| GC36429084 | Y2_Tnp\|Zn_Tnp_IS91 | Mobile genetic elements |
| GC37976755 | rve | Mobile genetic elements |
| GC38073324 | Resolvase | Mobile genetic elements |
| **GC36641378** | | |
| GC1127895 | HTH_1\|LysR_substrate | |
| GC11699548 | adh_short_C2 | |
| GC14713096 | HTH_38\|rve | Mobile genetic elements |
| GC15813830 | DEDD_Tnp_IS110\|Transposase_20 | Mobile genetic elements |
| GC16001823 | HTH_1\|LysR_substrate | |
| GC16204486 | Resolvase | Mobile genetic elements |
| GC18411621 | DDE_Tnp_1_6\|DUF772 | Mobile genetic elements |
| GC23019517 | DDE_Tnp_1_4 | Mobile genetic elements |
| GC23227492 | GIIM\|RVT_1 | Mobile genetic elements |
| GC24192253 | HTH_32 | |
| GC25753333 | RVT_1 | Mobile genetic elements |
| GC2923309 | DDE_Tnp_1\|DUF772 | Mobile genetic elements |
| GC30561759 | Phage_integrase\|Phage_int_SAM_4 | Mobile genetic elements |
| GC30764448 | DDE_Tnp_1 | Mobile genetic elements |
| GC33897398 | Transposase_20 | Mobile genetic elements |
| GC35526541 | HTH\|HTH | |
| GC36071194 | DDE_3 | Mobile genetic elements |
| GC36115360 | DDE_Tnp_IS240 | Mobile genetic elements |
| GC36597656 | Recombinase\|Resolvase | Mobile genetic elements |
| GC36768812 | IstB_IS21 | Mobile genetic elements |
| GC37209285 | DDE\|HTH | Mobile genetic elements |
| GC37272480 | DDE\|HTH | Mobile genetic elements |
| GC37552624 | Recombinase\|Resolvase | Mobile genetic elements |
| GC38232761 | DDE_3 | Mobile genetic elements |
| GC38440389 | Y2_Tnp\|Zn_Tnp_IS91 | Mobile genetic elements |
| GC38580544 | Transposase_20 | Mobile genetic elements |
| GC465251 | ACR_tran\|ASL_C2 | |
| GC5059750 | Y2_Tnp\|Zn_Tnp_IS91 | Mobile genetic elements |
| GC8673204 | ACR_tran | |
| GC9296586 | DDE_3\|HTH_32 | Mobile genetic elements |
| GC1011580 | Recombinase\|Resolvase | Mobile genetic elements |
| GC11494356 | GIIM\|RVT_1 | Mobile genetic elements |
| GC12214551 | HTH_Tnp_1 | Mobile genetic elements |
| GC13014417 | Recombinase\|Resolvase | Mobile genetic elements |
| GC1804679 | Phage_integrase | Mobile genetic elements |
| GC19517857 | rve_3\|rve | Mobile genetic elements |
| GC21347156 | GIIM\|RVT_1 | Mobile genetic elements |
| GC24933148 | DDE_Tnp_IS1595\|Zn_Tnp_IS1595 | Mobile genetic elements |
| GC25354868 | HTH_32\|HTH_33 | |
| GC27144963 | Transposase_20 | Mobile genetic elements |
| GC31612606 | GIIM\|RVT_1 | Mobile genetic elements |
| GC31969864 | DDE_Tnp_1\|DUF772 | Mobile genetic elements |
| GC31986313 | DEDD_Tnp_IS110\|Transposase_20 | Mobile genetic elements |
| GC33500111 | DEDD_Tnp_IS110\|Transposase | Mobile genetic elements |
| GC35554125 | DEDD_Tnp_IS110\|Transposase | Mobile genetic elements |
| GC35713887 | HTH_Tnp_1 | Mobile genetic elements |
| GC36248678 | GIIM\|RVT | Mobile genetic elements |
| GC36364646 | DDE_Tnp\|DUF4372 | Mobile genetic elements |
| GC36543908 | DDE_5 | Mobile genetic elements |
| GC37678183 | DDE_Tnp_1_6 | Mobile genetic elements |
| GC38000184 | HTH_Tnp\|TSC22 | Mobile genetic elements |
| GC39015032 | DEDD_Tnp_IS110\|Transposase | Mobile genetic elements |
| GC39316024 | GIIM\|RVT | Mobile genetic elements |
| GC39562619 | Recombinase\|Resolvase | Mobile genetic elements |
| GC39642572 | GIIM\|RVT | Mobile genetic elements |
| GC39817105 | Phage_integrase | Mobile genetic elements |
| GC39878062 | ACR_tran | |
| GC39929056 | Y2_Tnp\|Zn_Tnp_IS91 | Mobile genetic elements |
| GC5906625 | DDE_3\|HTH_32 | Mobile genetic elements |
| GC26926321 | DDE_Tnp_1_2 | Mobile genetic elements |
| GC27189918 | Y2_Tnp | Mobile genetic elements |
| GC35916463 | TnpB_IS66 | Mobile genetic elements |
| GC36429084 | Y2_Tnp\|Zn_Tnp_IS91 | Mobile genetic elements |
| GC37976755 | rve | Mobile genetic elements |
| GC38073324 | Resolvase | Mobile genetic elements |
| **GC39021886** | | |
| GC15813830 | DEDD_Tnp_IS110\|Transposase_20 | Mobile genetic elements |
| GC23019517 | DDE_Tnp_1_4 | Mobile genetic elements |
| GC25008120 | DEDD_Tnp_IS110\|Transposase_20 | Mobile genetic elements |
| GC25753333 | RVT_1 | Mobile genetic elements |
| GC30561759 | Phage_integrase\|Phage_int_SAM_4 | Mobile genetic elements |
| GC36071194 | DDE_3 | Mobile genetic elements |
| GC37208808 | RVT_1 | Mobile genetic elements |
| GC37209285 | DDE\|HTH | Mobile genetic elements |
| GC37272480 | DDE\|HTH | Mobile genetic elements |
| GC38440389 | Y2_Tnp\|Zn_Tnp_IS91 | Mobile genetic elements |
| GC38580544 | Transposase_20 | Mobile genetic elements |
| GC5059750 | Y2_Tnp\|Zn_Tnp_IS91 | Mobile genetic elements |
| GC15453272 | HTH_29\|rve_3\|rve | Mobile genetic elements |
| GC19517857 | rve_3\|rve | Mobile genetic elements |
| GC24933148 | DDE_Tnp_IS1595\|Zn_Tnp_IS1595 | Mobile genetic elements |
| GC25354868 | HTH_32\|HTH_33 | |
| GC31969864 | DDE_Tnp_1\|DUF772 | Mobile genetic elements |
| GC31986313 | DEDD_Tnp_IS110\|Transposase_20 | Mobile genetic elements |
| GC33500111 | DEDD_Tnp_IS110\|Transposase | Mobile genetic elements |
| GC36364646 | DDE_Tnp\|DUF4372 | Mobile genetic elements |
| GC38000184 | HTH_Tnp\|TSC22 | Mobile genetic elements |
| GC39316024 | GIIM\|RVT | Mobile genetic elements |
| GC27189918 | Y2_Tnp | Mobile genetic elements |
| **GC39495362** | | |
| GC11144186 | Transposase_mut | Mobile genetic elements |
| GC30561759 | Phage_integrase\|Phage_int_SAM_4 | Mobile genetic elements |
| GC36071194 | DDE_3 | Mobile genetic elements |
| GC36115360 | DDE_Tnp_IS240 | Mobile genetic elements |
| GC37208808 | RVT_1 | Mobile genetic elements |
| GC37272480 | DDE\|HTH | Mobile genetic elements |
| GC38440389 | Y2_Tnp\|Zn_Tnp_IS91 | Mobile genetic elements |
| GC5059750 | Y2_Tnp\|Zn_Tnp_IS91 | Mobile genetic elements |
| GC25354868 | HTH_32\|HTH_33 | |
| GC31969864 | DDE_Tnp_1\|DUF772 | Mobile genetic elements |
| GC36364646 | DDE_Tnp\|DUF4372 | Mobile genetic elements |
| GC38000184 | HTH_Tnp\|TSC22 | Mobile genetic elements |
| GC39015032 | DEDD_Tnp_IS110\|Transposase | Mobile genetic elements |
| GC39642572 | GIIM\|RVT | Mobile genetic elements |
| GC38392731 | TnpB_IS66 | Mobile genetic elements |
| **GC31831041** | | |
| GC23019517 | DDE_Tnp_1_4 | Mobile genetic elements |
| GC34251202 | DDE_Tnp_ISAZ013 | Mobile genetic elements |
| **GC31713130** | | |
| GC16107005 | DDE_5\|DDE_Tnp_1 | Mobile genetic elements |
| **GC34767203** | | |
| GC11699548 | adh_short_C2 | |
| GC11967938 | adh_short_C2\|POR | |
| GC15813830 | DEDD_Tnp_IS110\|Transposase_20 | Mobile genetic elements |
| GC16001823 | HTH_1\|LysR_substrate | |
| GC18411621 | DDE_Tnp_1_6\|DUF772 | Mobile genetic elements |
| GC19825056 | GMC_oxred_C\|GMC_oxred_N | |
| GC23019517 | DDE_Tnp_1_4 | Mobile genetic elements |
| GC23227492 | GIIM\|RVT_1 | Mobile genetic elements |
| GC24192253 | HTH_32 | |
| GC25753333 | RVT_1 | Mobile genetic elements |
| GC2923309 | DDE_Tnp_1\|DUF772 | Mobile genetic elements |
| GC30561759 | Phage_integrase\|Phage_int_SAM_4 | Mobile genetic elements |
| GC30764448 | DDE_Tnp_1 | Mobile genetic elements |
| GC32620141 | DDE_Tnp_1_4 | Mobile genetic elements |
| GC33897398 | Transposase_20 | Mobile genetic elements |
| GC34651936 | DDE_Tnp_1 | Mobile genetic elements |
| GC36115360 | DDE_Tnp_IS240 | Mobile genetic elements |
| GC36597656 | Recombinase\|Resolvase | Mobile genetic elements |
| GC36768812 | IstB_IS21 | Mobile genetic elements |
| GC37272480 | DDE\|HTH | Mobile genetic elements |
| GC38220472 | GIIM\|RVT | Mobile genetic elements |
| GC38440389 | Y2_Tnp\|Zn_Tnp_IS91 | Mobile genetic elements |
| GC38580544 | Transposase_20 | Mobile genetic elements |
| GC4229464 | GMC_oxred_N | |
| GC5059750 | Y2_Tnp\|Zn_Tnp_IS91 | Mobile genetic elements |
| GC8673204 | ACR_tran | |
| GC9296586 | DDE_3\|HTH_32 | Mobile genetic elements |
| GC1011580 | Recombinase\|Resolvase | Mobile genetic elements |
| GC12214551 | HTH_Tnp_1 | Mobile genetic elements |
| GC13014417 | Recombinase\|Resolvase | Mobile genetic elements |
| GC15453272 | HTH_29\|rve_3\|rve | Mobile genetic elements |
| GC19517857 | rve_3\|rve | Mobile genetic elements |
| GC24933148 | DDE_Tnp_IS1595\|Zn_Tnp_IS1595 | Mobile genetic elements |
| GC31612606 | GIIM\|RVT_1 | Mobile genetic elements |
| GC31969864 | DDE_Tnp_1\|DUF772 | Mobile genetic elements |
| GC35554125 | DEDD_Tnp_IS110\|Transposase | Mobile genetic elements |
| GC35554439 | Transposase_mut | Mobile genetic elements |
| GC35713887 | HTH_Tnp_1 | Mobile genetic elements |
| GC36248678 | GIIM\|RVT | Mobile genetic elements |
| GC36364646 | DDE_Tnp\|DUF4372 | Mobile genetic elements |
| GC39316024 | GIIM\|RVT | Mobile genetic elements |
| GC39562619 | Recombinase\|Resolvase | Mobile genetic elements |
| GC39642572 | GIIM\|RVT | Mobile genetic elements |
| GC39817105 | Phage_integrase | Mobile genetic elements |
| GC39878062 | ACR_tran | |
| GC39929056 | Y2_Tnp\|Zn_Tnp_IS91 | Mobile genetic elements |
| GC5906625 | DDE_3\|HTH_32 | Mobile genetic elements |
| GC26926321 | DDE_Tnp_1_2 | Mobile genetic elements |
| GC27189918 | Y2_Tnp | Mobile genetic elements |
| GC35916463 | TnpB_IS66 | Mobile genetic elements |
| GC36429084 | Y2_Tnp\|Zn_Tnp_IS91 | Mobile genetic elements |
| GC37236299 | Y2_Tnp\|Zn_Tnp_IS91 | Mobile genetic elements |
| GC37345069 | DDE_Tnp\|DUF772 | Mobile genetic elements |
| GC37976755 | rve | Mobile genetic elements |
| GC38073324 | Resolvase | Mobile genetic elements |
| GC25129005 | DDE_Tnp_1\|DUF4096 | Mobile genetic elements |
| **GC35697170** | | |
| GC14713096 | HTH_38\|rve | Mobile genetic elements |
| GC15813830 | DEDD_Tnp_IS110\|Transposase_20 | Mobile genetic elements |
| GC18411621 | DDE_Tnp_1_6\|DUF772 | Mobile genetic elements |
| GC23019517 | DDE_Tnp_1_4 | Mobile genetic elements |
| GC23227492 | GIIM\|RVT_1 | Mobile genetic elements |
| GC24192253 | HTH_32 | |
| GC25008120 | DEDD_Tnp_IS110\|Transposase_20 | Mobile genetic elements |
| GC25753333 | RVT_1 | Mobile genetic elements |
| GC32367139 | DEDD_Tnp_IS110\|Transposase_20 | Mobile genetic elements |
| GC34251202 | DDE_Tnp_ISAZ013 | Mobile genetic elements |
| GC36558232 | Transposase_mut | Mobile genetic elements |
| GC37209285 | DDE\|HTH | Mobile genetic elements |
| GC38440389 | Y2_Tnp\|Zn_Tnp_IS91 | Mobile genetic elements |
| GC38580544 | Transposase_20 | Mobile genetic elements |
| GC12214551 | HTH_Tnp_1 | Mobile genetic elements |
| GC19517857 | rve_3\|rve | Mobile genetic elements |
| GC21347156 | GIIM\|RVT_1 | Mobile genetic elements |
| GC27144963 | Transposase_20 | Mobile genetic elements |
| GC31969864 | DDE_Tnp_1\|DUF772 | Mobile genetic elements |
| GC31986313 | DEDD_Tnp_IS110\|Transposase_20 | Mobile genetic elements |
| GC33500111 | DEDD_Tnp_IS110\|Transposase | Mobile genetic elements |
| GC35554125 | DEDD_Tnp_IS110\|Transposase | Mobile genetic elements |
| GC38000184 | HTH_Tnp\|TSC22 | Mobile genetic elements |
| GC39015032 | DEDD_Tnp_IS110\|Transposase | Mobile genetic elements |
| GC39316024 | GIIM\|RVT | Mobile genetic elements |
| GC41300537 | DDE_5 | Mobile genetic elements |
| GC5906625 | DDE_3\|HTH_32 | Mobile genetic elements |
| GC10026933 | DEDD_Tnp_IS110 | Mobile genetic elements |
| GC27189918 | Y2_Tnp | Mobile genetic elements |
| GC32855953 | Transposase_mut | Mobile genetic elements |
| GC36429084 | Y2_Tnp\|Zn_Tnp_IS91 | Mobile genetic elements |
| GC37976755 | rve | Mobile genetic elements |
| GC38073324 | Resolvase | Mobile genetic elements |
| GC25129005 | DDE_Tnp_1\|DUF4096 | Mobile genetic elements |
| **GC30406655** | | |
| GC11699548 | adh_short_C2 | |
| GC16001823 | HTH_1\|LysR_substrate | |
| GC18411621 | DDE_Tnp_1_6\|DUF772 | Mobile genetic elements |
| GC30561759 | Phage_integrase\|Phage_int_SAM_4 | Mobile genetic elements |
| GC33897398 | Transposase_20 | Mobile genetic elements |
| GC36768812 | IstB_IS21 | Mobile genetic elements |
| GC37272480 | DDE\|HTH | Mobile genetic elements |
| GC38440389 | Y2_Tnp\|Zn_Tnp_IS91 | Mobile genetic elements |
| GC38580544 | Transposase_20 | Mobile genetic elements |
| GC5059750 | Y2_Tnp\|Zn_Tnp_IS91 | Mobile genetic elements |
| GC1011580 | Recombinase\|Resolvase | Mobile genetic elements |
| GC13014417 | Recombinase\|Resolvase | Mobile genetic elements |
| GC19517857 | rve_3\|rve | Mobile genetic elements |
| GC31969864 | DDE_Tnp_1\|DUF772 | Mobile genetic elements |
| GC36248678 | GIIM\|RVT | Mobile genetic elements |
| GC37678183 | DDE_Tnp_1_6 | Mobile genetic elements |
| GC39015032 | DEDD_Tnp_IS110\|Transposase | Mobile genetic elements |
| GC39562619 | Recombinase\|Resolvase | Mobile genetic elements |
| GC39878062 | ACR_tran | |
| GC39929056 | Y2_Tnp\|Zn_Tnp_IS91 | Mobile genetic elements |
| GC9076628 | DDE_Tnp_1\|DUF772 | Mobile genetic elements |
| GC26926321 | DDE_Tnp_1_2 | Mobile genetic elements |
| GC27189918 | Y2_Tnp | Mobile genetic elements |
| GC35916463 | TnpB_IS66 | Mobile genetic elements |
| GC37976755 | rve | Mobile genetic elements |
| GC38073324 | Resolvase | Mobile genetic elements |
| **GC37353006** | | |
| GC27386295 | HTH_21\|rve | Mobile genetic elements |
| GC37377483 | HTH\|rve | Mobile genetic elements |
| **GC36001967** | | |
| GC23019517 | DDE_Tnp_1_4 | Mobile genetic elements |
| GC24192253 | HTH_32 | |
| GC25753333 | RVT_1 | Mobile genetic elements |
| GC30561759 | Phage_integrase\|Phage_int_SAM_4 | Mobile genetic elements |
| GC36768812 | IstB_IS21 | Mobile genetic elements |
| GC37552624 | Recombinase\|Resolvase | Mobile genetic elements |
| GC39056764 | IstB_IS21 | Mobile genetic elements |
| GC5059750 | Y2_Tnp\|Zn_Tnp_IS91 | Mobile genetic elements |
| GC13014417 | Recombinase\|Resolvase | Mobile genetic elements |
| GC21347156 | GIIM\|RVT_1 | Mobile genetic elements |
| GC24933148 | DDE_Tnp_IS1595\|Zn_Tnp_IS1595 | Mobile genetic elements |
| GC36364646 | DDE_Tnp\|DUF4372 | Mobile genetic elements |
| GC39316024 | GIIM\|RVT | Mobile genetic elements |
| GC39562619 | Recombinase\|Resolvase | Mobile genetic elements |
| GC5906625 | DDE_3\|HTH_32 | Mobile genetic elements |
| GC26787771 | RVT_1 | Mobile genetic elements |
| GC37345069 | DDE_Tnp\|DUF772 | Mobile genetic elements |
| GC37976755 | rve | Mobile genetic elements |
| **GC39377992** | | |
| GC15813830 | DEDD_Tnp_IS110\|Transposase_20 | Mobile genetic elements |
| GC18411621 | DDE_Tnp_1_6\|DUF772 | Mobile genetic elements |
| GC23019517 | DDE_Tnp_1_4 | Mobile genetic elements |
| GC23227492 | GIIM\|RVT_1 | Mobile genetic elements |
| GC2923309 | DDE_Tnp_1\|DUF772 | Mobile genetic elements |
| GC30561759 | Phage_integrase\|Phage_int_SAM_4 | Mobile genetic elements |
| GC33897398 | Transposase_20 | Mobile genetic elements |
| GC35833714 | rve\|rve | Mobile genetic elements |
| GC15453272 | HTH_29\|rve_3\|rve | Mobile genetic elements |
| GC19517857 | rve_3\|rve | Mobile genetic elements |
| GC36364646 | DDE_Tnp\|DUF4372 | Mobile genetic elements |
| GC39316024 | GIIM\|RVT | Mobile genetic elements |
| GC39929056 | Y2_Tnp\|Zn_Tnp_IS91 | Mobile genetic elements |
| GC5906625 | DDE_3\|HTH_32 | Mobile genetic elements |
| GC10026933 | DEDD_Tnp_IS110 | Mobile genetic elements |
| GC27189918 | Y2_Tnp | Mobile genetic elements |
| GC37236299 | Y2_Tnp\|Zn_Tnp_IS91 | Mobile genetic elements |
| **GC37458337** | | |
| GC25753333 | RVT_1 | Mobile genetic elements |
| **GC36965309** | | |
| GC11699548 | adh_short_C2 | |
| GC11967938 | adh_short_C2\|POR | |
| GC15813830 | DEDD_Tnp_IS110\|Transposase_20 | Mobile genetic elements |
| GC16001823 | HTH_1\|LysR_substrate | |
| GC17942492 | BON\|BON\|BON | |
| GC18411621 | DDE_Tnp_1_6\|DUF772 | Mobile genetic elements |
| GC19108944 | F420_oxidored | |
| GC19825056 | GMC_oxred_C\|GMC_oxred_N | |
| GC20124390 | ACT\|GMC_oxred_N | |
| GC23019517 | DDE_Tnp_1_4 | Mobile genetic elements |
| GC23227492 | GIIM\|RVT_1 | Mobile genetic elements |
| GC24192253 | HTH_32 | |
| GC25753333 | RVT_1 | Mobile genetic elements |
| GC30561759 | Phage_integrase\|Phage_int_SAM_4 | Mobile genetic elements |
| GC30764448 | DDE_Tnp_1 | Mobile genetic elements |
| GC31902049 | ACR_tran | |
| GC33897398 | Transposase_20 | Mobile genetic elements |
| GC35526541 | HTH\|HTH | |
| GC36115360 | DDE_Tnp_IS240 | Mobile genetic elements |
| GC36597656 | Recombinase\|Resolvase | Mobile genetic elements |
| GC36768812 | IstB_IS21 | Mobile genetic elements |
| GC37209285 | DDE\|HTH | Mobile genetic elements |
| GC37272480 | DDE\|HTH | Mobile genetic elements |
| GC37499516 | RVT_1 | Mobile genetic elements |
| GC38440389 | Y2_Tnp\|Zn_Tnp_IS91 | Mobile genetic elements |
| GC38580544 | Transposase_20 | Mobile genetic elements |
| GC465251 | ACR_tran\|ASL_C2 | |
| GC5059750 | Y2_Tnp\|Zn_Tnp_IS91 | Mobile genetic elements |
| GC8673204 | ACR_tran | |
| GC1011580 | Recombinase\|Resolvase | Mobile genetic elements |
| GC13014417 | Recombinase\|Resolvase | Mobile genetic elements |
| GC1804679 | Phage_integrase | Mobile genetic elements |
| GC19517857 | rve_3\|rve | Mobile genetic elements |
| GC24933148 | DDE_Tnp_IS1595\|Zn_Tnp_IS1595 | Mobile genetic elements |
| GC27144963 | Transposase_20 | Mobile genetic elements |
| GC31612606 | GIIM\|RVT_1 | Mobile genetic elements |
| GC31969864 | DDE_Tnp_1\|DUF772 | Mobile genetic elements |
| GC31986313 | DEDD_Tnp_IS110\|Transposase_20 | Mobile genetic elements |
| GC35554125 | DEDD_Tnp_IS110\|Transposase | Mobile genetic elements |
| GC35713887 | HTH_Tnp_1 | Mobile genetic elements |
| GC36110782 | DEDD_Tnp_IS110\|Transposase | Mobile genetic elements |
| GC36248678 | GIIM\|RVT | Mobile genetic elements |
| GC36364646 | DDE_Tnp\|DUF4372 | Mobile genetic elements |
| GC37678183 | DDE_Tnp_1_6 | Mobile genetic elements |
| GC38000184 | HTH_Tnp\|TSC22 | Mobile genetic elements |
| GC39015032 | DEDD_Tnp_IS110\|Transposase | Mobile genetic elements |
| GC39316024 | GIIM\|RVT | Mobile genetic elements |
| GC39562619 | Recombinase\|Resolvase | Mobile genetic elements |
| GC39642572 | GIIM\|RVT | Mobile genetic elements |
| GC39817105 | Phage_integrase | Mobile genetic elements |
| GC39878062 | ACR_tran | |
| GC39929056 | Y2_Tnp\|Zn_Tnp_IS91 | Mobile genetic elements |
| GC5906625 | DDE_3\|HTH_32 | Mobile genetic elements |
| GC9076628 | DDE_Tnp_1\|DUF772 | Mobile genetic elements |
| GC10026933 | DEDD_Tnp_IS110 | Mobile genetic elements |
| GC26926321 | DDE_Tnp_1_2 | Mobile genetic elements |
| GC27189918 | Y2_Tnp | Mobile genetic elements |
| GC35916463 | TnpB_IS66 | Mobile genetic elements |
| GC36429084 | Y2_Tnp\|Zn_Tnp_IS91 | Mobile genetic elements |
| GC37236299 | Y2_Tnp\|Zn_Tnp_IS91 | Mobile genetic elements |
| GC37345069 | DDE_Tnp\|DUF772 | Mobile genetic elements |
| GC37976755 | rve | Mobile genetic elements |
| GC38073324 | Resolvase | Mobile genetic elements |
| GC21618014 | GIIM\|RVT_1 | Mobile genetic elements |
| **GC33801135** | | |
| GC23227492 | GIIM\|RVT_1 | Mobile genetic elements |
| GC11494356 | GIIM\|RVT_1 | Mobile genetic elements |
| GC15453272 | HTH_29\|rve_3\|rve | Mobile genetic elements |
| GC24933148 | DDE_Tnp_IS1595\|Zn_Tnp_IS1595 | Mobile genetic elements |
| GC5906625 | DDE_3\|HTH_32 | Mobile genetic elements |
| GC27189918 | Y2_Tnp | Mobile genetic elements |
| **GC36701415** | | |
| GC11144186 | Transposase_mut | Mobile genetic elements |
| GC14713096 | HTH_38\|rve | Mobile genetic elements |
| GC15813830 | DEDD_Tnp_IS110\|Transposase_20 | Mobile genetic elements |
| GC16204486 | Resolvase | Mobile genetic elements |
| GC18411621 | DDE_Tnp_1_6\|DUF772 | Mobile genetic elements |
| GC23019517 | DDE_Tnp_1_4 | Mobile genetic elements |
| GC24192253 | HTH_32 | |
| GC25008120 | DEDD_Tnp_IS110\|Transposase_20 | Mobile genetic elements |
| GC25753333 | RVT_1 | Mobile genetic elements |
| GC30561759 | Phage_integrase\|Phage_int_SAM_4 | Mobile genetic elements |
| GC30764448 | DDE_Tnp_1 | Mobile genetic elements |
| GC32367139 | DEDD_Tnp_IS110\|Transposase_20 | Mobile genetic elements |
| GC33897398 | Transposase_20 | Mobile genetic elements |
| GC36071194 | DDE_3 | Mobile genetic elements |
| GC36597656 | Recombinase\|Resolvase | Mobile genetic elements |
| GC37208808 | RVT_1 | Mobile genetic elements |
| GC37209285 | DDE\|HTH | Mobile genetic elements |
| GC37499516 | RVT_1 | Mobile genetic elements |
| GC38440389 | Y2_Tnp\|Zn_Tnp_IS91 | Mobile genetic elements |
| GC5059750 | Y2_Tnp\|Zn_Tnp_IS91 | Mobile genetic elements |
| GC12214551 | HTH_Tnp_1 | Mobile genetic elements |
| GC15453272 | HTH_29\|rve_3\|rve | Mobile genetic elements |
| GC1804679 | Phage_integrase | Mobile genetic elements |
| GC19517857 | rve_3\|rve | Mobile genetic elements |
| GC21347156 | GIIM\|RVT_1 | Mobile genetic elements |
| GC24933148 | DDE_Tnp_IS1595\|Zn_Tnp_IS1595 | Mobile genetic elements |
| GC31969864 | DDE_Tnp_1\|DUF772 | Mobile genetic elements |
| GC31986313 | DEDD_Tnp_IS110\|Transposase_20 | Mobile genetic elements |
| GC35554125 | DEDD_Tnp_IS110\|Transposase | Mobile genetic elements |
| GC35713887 | HTH_Tnp_1 | Mobile genetic elements |
| GC36110782 | DEDD_Tnp_IS110\|Transposase | Mobile genetic elements |
| GC36364646 | DDE_Tnp\|DUF4372 | Mobile genetic elements |
| GC38000184 | HTH_Tnp\|TSC22 | Mobile genetic elements |
| GC39015032 | DEDD_Tnp_IS110\|Transposase | Mobile genetic elements |
| GC39316024 | GIIM\|RVT | Mobile genetic elements |
| GC39562619 | Recombinase\|Resolvase | Mobile genetic elements |
| GC39642572 | GIIM\|RVT | Mobile genetic elements |
| GC39929056 | Y2_Tnp\|Zn_Tnp_IS91 | Mobile genetic elements |
| GC5906625 | DDE_3\|HTH_32 | Mobile genetic elements |
| GC9076628 | DDE_Tnp_1\|DUF772 | Mobile genetic elements |
| GC10026933 | DEDD_Tnp_IS110 | Mobile genetic elements |
| GC26787771 | RVT_1 | Mobile genetic elements |
| GC26926321 | DDE_Tnp_1_2 | Mobile genetic elements |
| GC27189918 | Y2_Tnp | Mobile genetic elements |
| GC33398825 | DDE_5 | Mobile genetic elements |
| GC36429084 | Y2_Tnp\|Zn_Tnp_IS91 | Mobile genetic elements |
| GC37976755 | rve | Mobile genetic elements |
| GC38073324 | Resolvase | Mobile genetic elements |
| GC21618014 | GIIM\|RVT_1 | Mobile genetic elements |
