## Supplementary figures and images for "An ecological perspective on microbial genes of unknown function in soil"

### Figure S5

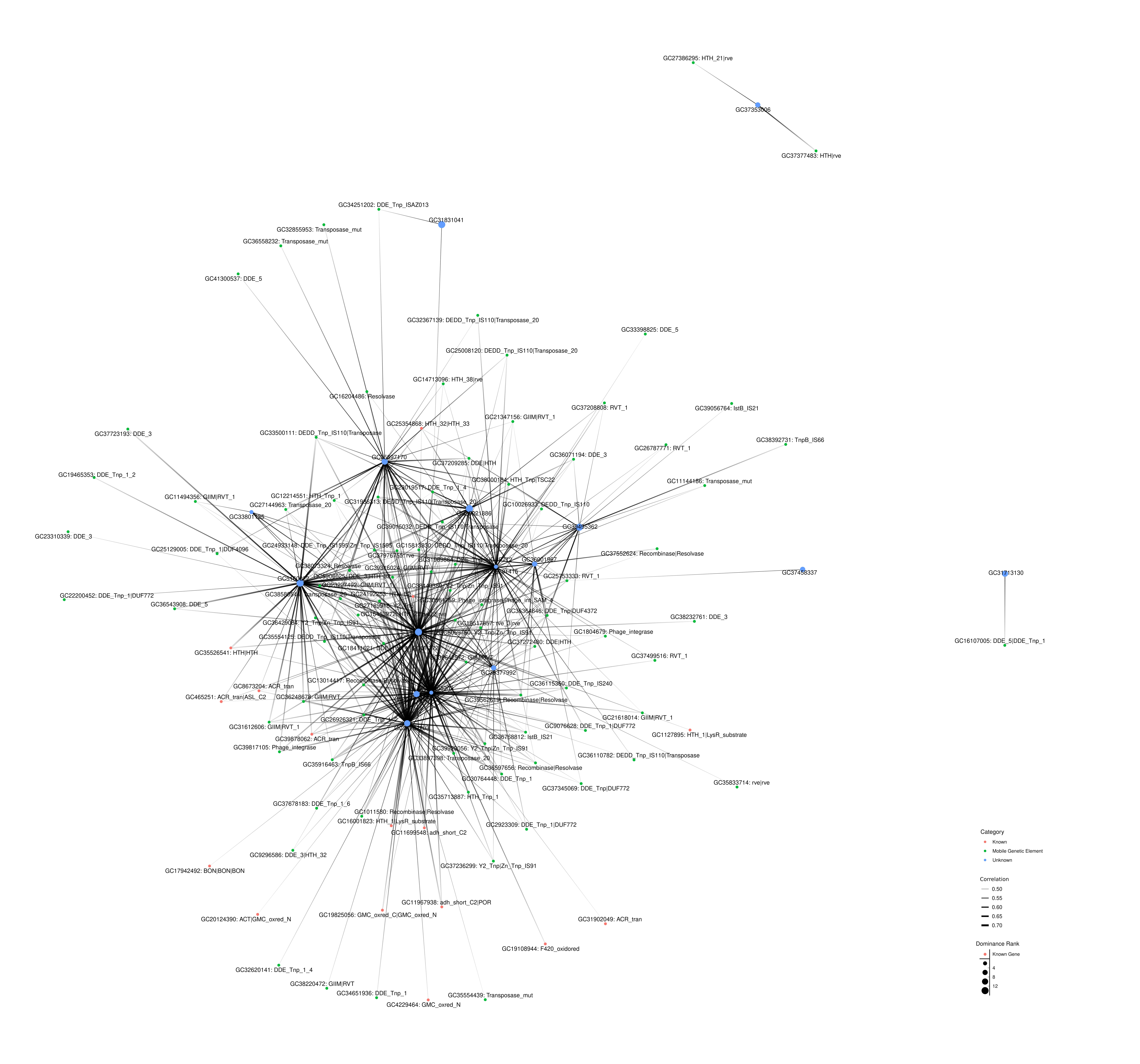
